# Coordinated behavioural and neural variability in hippocampal reward coding

**DOI:** 10.64898/2026.08.13.744587

**Authors:** Charline Tessereau, Peter Dayan

## Abstract

An environment can be interpreted in multiple ways, with implications for behaviour and neural representations and for revision in these when circumstances change. Using two-photon calcium imaging in mouse CA1 during navigation under structured reward-location uncertainty, we found correlations between variability in anticipatory behaviour and the organization of place fields with respect to position and reward. Greater spatial anticipation was associated with enhanced reward-centred coding and increased generalisability of reward-aligned representations.

---

Consider a visually stable environment in which reward occurs at locations that vary stochastically. Stable track position and reward location on a given traversal need not coincide, and animals may weight these sources of structure differently, such that the same sensory environment supports distinct behavioural strategies and neural representations [1, 2]. Characterising this variation is important because natural environments are often stable in some respects but uncertain in others.

Activity in hippocampal area CA1 famously covaries with spatial position [3], but is also shaped by visual context [4, 5], reward and goals [6, 7], task structure [8–10], and internal state [11]. When reward moves within an otherwise unchanged environment, some place fields remain fixed in absolute position, whereas others maintain their alignment to reward as it shifts. We asked whether stable individual differences in spatial reward anticipation predict the balance between these forms of coding.

To test this, we used a virtual-reality stochastic reward navigation task [10]. Nine head-fixed mice ran along a 3-m virtual linear track while licking was monitored and dorsal CA1 pyramidal cells expressing jGCaMP8m were recorded using two-photon calcium imaging (Fig. 1A–C). During training and the first part of a ‘switch’ session, one water reward was delivered on each traversal at one of ten uniformly sampled locations spanning a 1m zone. In the switch session, without warning, the reward distribution then shifted to two locations spanning a 10cm zone further down the visual track (Fig. 1D).

**Fig. 1.**
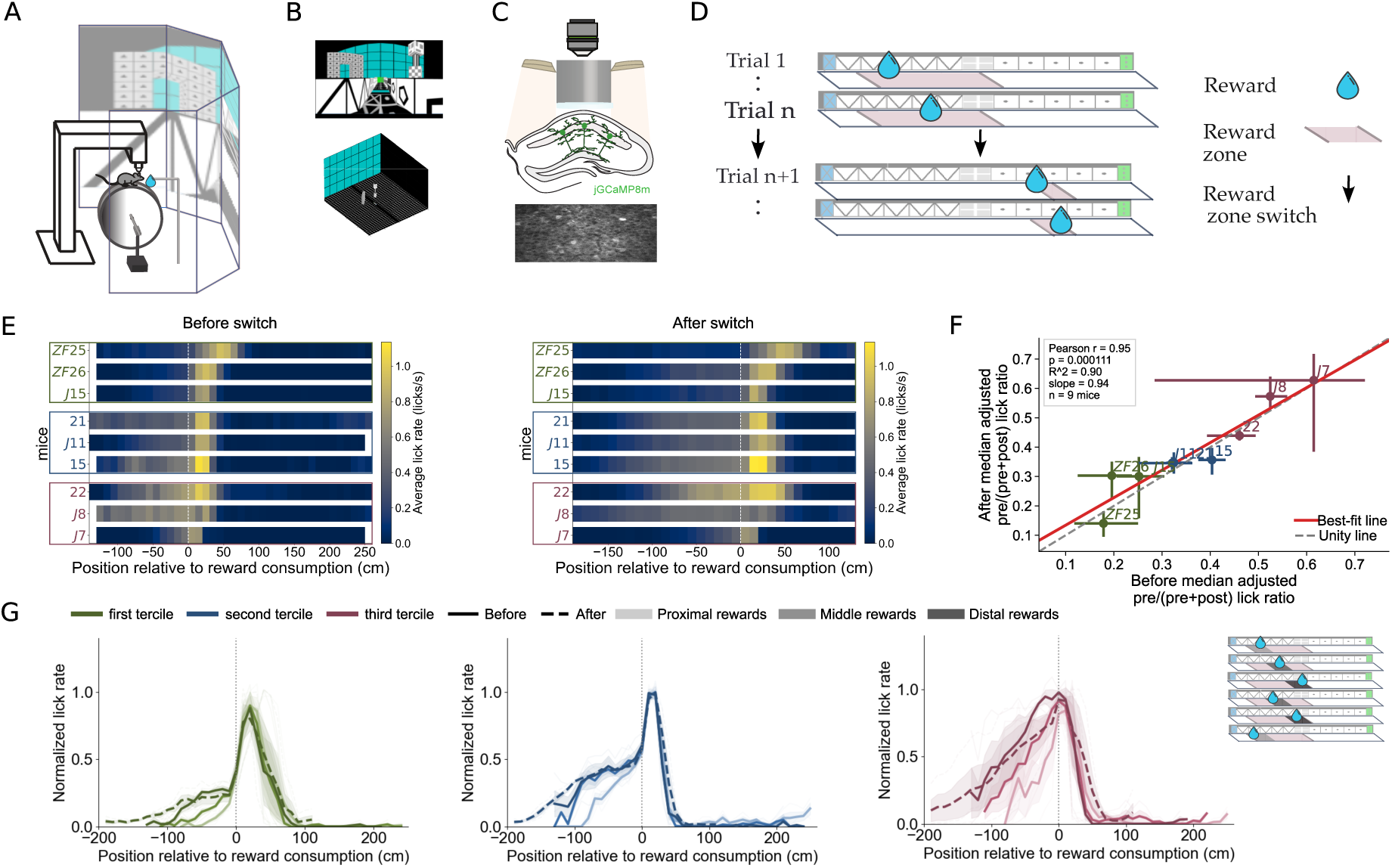
Mice adopt distinct reward-responsive and anticipatory licking strategies under reward-location uncertainty. **A:** Schematic of the virtual reality (VR) apparatus: the licking behaviour of mice was recorded as they ran on a wheel with a closed-loop connection to the visual movement of a virtual track on the display. When the mice reached the end of the track, the screen went black for 3 seconds and mice were teleported to the start of the track for the next run. **B:** Visualization of the track used for VR in this paper. Top: front view of the track. Bottom: 3D view of the track, showing the relative perspective with distal cues. **C:** Schematic of two-photon calcium imaging of mouse CA1 neurons (green colours) expressing jGCaMP8m (top) and imaging field of view (bottom). **D:** Schematic of task design. Mice ran on a 1D virtual reality environment in which a water reward is initially delivered each trial within a 1m reward zone once per traversal of a 3m long linear track (and could subsequently be consumed anywhere by licking). During training and at the start of the switch session considered in this paper, on every run, the reward location was randomly drawn uniformly from one of 10 potential locations evenly spaced within the 1m wide zone. During the session, on trial *n* + 1 (*n* = 36.8 *±* 4.5 std), without prior notice, the reward delivery location switched to one of two positions at the edges of a more remote 10cm zone. Additional task and training details in Methods. **E:** Left: Average lick rate aligned to reward consumption before the switch for each of the nine mice. Right: Corresponding reward-consumption-aligned average after the switch. Reward consumption, defined as the first lick after reward delivery, is located at 0 cm. Mice are ordered from the most reward-responsive (top) to the most spatially anticipatory (bottom). The colour of mice labels and highlighted in a rectangle indicates terciles: green is used for most reward-responsive tercile (top), blue for intermediate tercile (middle), and purple for the most anticipatory tercile (bottom). **F:** Relationship between each mouse’s median adjusted pre/(pre+post) lick ratio before (*x* axis) vs. after the switch (*y* axis). Each point is one mouse; horizontal and vertical bars show the interquartile range across values for that mouse before and after the switch, respectively. Label colour indicates the terciles of the animals: green for the first low-anticipatory, reward-responsive, tercile; blue for the middle tercile, purple for the high anticipatory tercile. The red line is the least-squares linear fit and the grey dashed line is the unity line. Pearson’s correlation was computed across mouse medians (*r* = 0.95, two-sided *p* = 1.11 *×* 10*^−^*^4^). The displayed slope (0.94) is the slope of the best-fit line for after-switch ratio as a function of before-switch ratio; *R*^2^ = 0.90 is the coefficient of determination. **G:** Reward-consumption-aligned lick rate for proximal reward trials (light shade), middle reward trials (intermediate shade) and distal reward trials (dark shade) before the switch (plain lines) and after the switch (dashed lines) for left/green: Lick rate for the most reward-responsive tercile of animals; middle/blue: the middle tercile animals, and right/purple: for the top tercile reward anticipatory animals. Inset on the top right portrays how trials are separated: to look at the effect of a continuously changing reward location before the switch, we separate the trials based on where the reward was consumed in the zone (see Methods, between proximal (light shades), middle (intermediary shades) and distal trials (dark shades).

All mice localised licking around the rewarded region (Fig. 1E), consistent with learning its distribution. Indeed, in the broad reward zone, all mice expressed spatially modulated pre-reward licking and licked more when reward was farther from the start of the track (’distal’) than near (’proximal’; one-sided Wilcoxon signed-rank test on mouse-level distal–proximal differences, *p* = 0.00195; Fig. 1G and Supplementary Fig. S1D for within-mice statistics), showing use of spatial information for anticipation. Reward-consumption-aligned licking profiles nevertheless differed markedly across animals (Fig. 1E and Supplementary Fig. S2). We quantified this behavioural variation using an anticipatory licking index, computed relative to reward delivery as the mean lick rate before delivery divided by the sum of matched preand post-delivery lick rates (Supplementary Fig. S1A and Methods). The index placed mice along a continuum from more reward-responsive profiles to more spatially anticipatory, with median indices ranging from 0.15 to 0.62 across mice. Tercile-averaged consumption-aligned profiles made the distinction explicit (although all inferential analyses were conducted at the mouse level): at the more reward-responsive end, spatially modulated pre-reward licking was followed by a dominant post-delivery bout, whereas at the more spatially anticipatory end, lick rate approached its peak before consumption (Fig. 1G). The behavioural ordering was stable across reward relocation (preversus post-switch *r* = 0.95, two-sided *p* = 1.11 *×* 10*^−^*^4^; Fig. 1F; dashed lines in Fig. 1G).

We next asked whether the behavioural continuum was associated with distinct patterns of CA1 coding when the reward location varied within the pre-switch broad zone. We compared proximal and distal reward-consumption trials, computing cross-validated maps in reward-consumption-aligned and absolute-position coordinates (Fig. 2A–C; see Methods). We classified cells as reward-aligned when their peaks fell within *−*20 to +30 cm of reward consumption on both trial types (yellow boxes in Fig. 2E–G), and as position-stable when their absolute-position peaks differed by no more than 30cm (yellow diagonal zone in Fig. S4E–G). The fraction of reward-aligned cells increased with the anticipatory licking index (*β* = 3.27; exact mouse-level permutation, two-sided *p*_perm_ = 0.0398; Fig. 2D–G). Position stability was not reliably associated with the anticipatory licking index in the primary continuous mouse-level analysis (exact mouse-label permutation test, two-sided *p*_perm_ = 0.3215; Supplementary Fig. S4), but showed a significant opposite trend across terciles (exact ordered-tercile permutation test, two-sided *p*_tercile_ = 0.040). All terciles also retained track-wide position coding (Supplementary Figs. S3 and S4). Representative cells illustrating both organisations are shown in Fig. 2I,J. Thus, stronger spatial anticipation was associated with more consistent reward-aligned coding across variable reward locations.

**Fig. 2.**
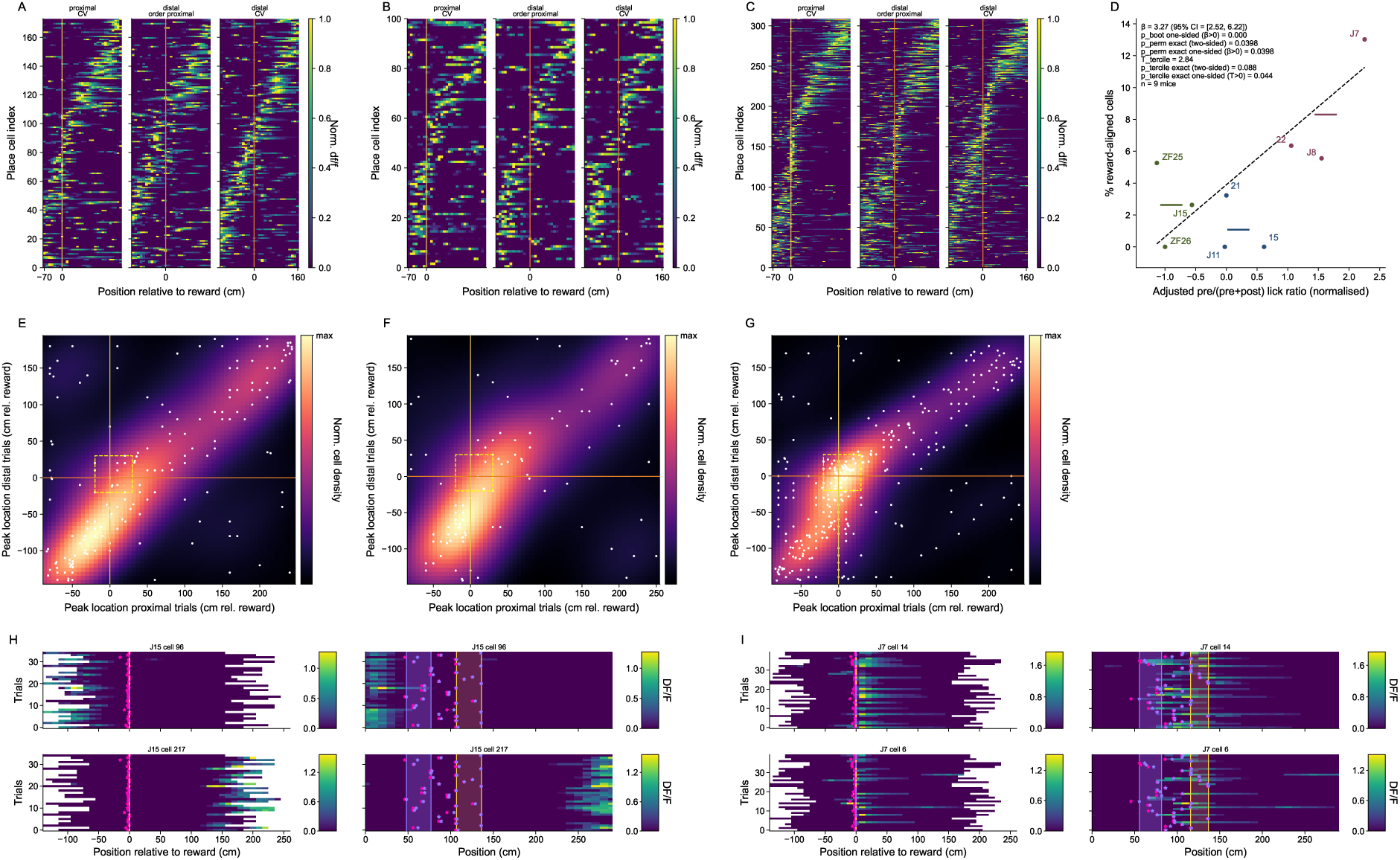
Spatial anticipatory licking predicts stronger reward-aligned CA1 coding across variable reward locations. **A:** Normalised cross-validated reward-consumption-aligned place maps for the first behavioural tercile, pooled across reward-responsive mice. Analyses use trials before the reward-location switch, divided according to reward-consumption location into proximal and distal reward trials. Each row represents a cell that met the spatial-information criterion for a significant place cell before the switch (see Methods). The left plot shows average activity on odd proximal trials, ordered by the peak of activity measured on even proximal trials; the middle plot shows activity on distal trials while retaining the proximal-trial ordering; and the right plot shows activity on odd distal trials reordered by the peak measured on even distal trials. Activity is aligned to reward consumption and normalised to each cell’s peak. Vertical lines mark reward consumption at 0 cm; yellow denotes proximal trials origin and orange denotes distal trials origin. **B, C:** Same as **A**, respectively, for mice in the second and third behavioural tercile, respectively. **D:** Across-mouse relationship between the normalised adjusted pre/(pre+post) lick ratio (*x* axis) and the proportion of reward-aligned cells across proximal and distal reward trials (*y* axis). The zone considered for reward-aligned cells is depicted using a yellow dashed rectangle in **E, F, G** and in supplementary S3D. Each point represents one mouse and is labelled by mouse identity; colours indicate lick-ratio terciles. Horizontal bars show the mean proportion of reward-aligned cells within each tercile. The dashed line shows the weighted linear-regression fit across mice (*β* = 3.27, 95% bootstrap CI [2.52, 6.22]; one-sided bootstrap test for *β >* 0, *p*_boot_ *<* 0.001; exact mouse-label permutation test: two-sided *p*_perm_ = 0.0398, one-sided *p*_perm_ = 0.0398). The ordered-tercile statistic was *T* = 2.84 (exact ordered-tercile permutation test: two-sided *p*_tercile_ = 0.088, one-sided *p*_tercile_ = 0.044; 1,680 ordered 3/3/3 assignments; *n* = 9 mice). **E:** Reward-aligned-frame proximal-versus-distal remapping for the same cells and mice as in **A**. The *x* and *y* coordinates show each cell’s peak reward-consumption-aligned activity location on proximal and distal trials, respectively. Each white point represents one place cell, and the magma heat map shows a normalised Gaussian kernel-density estimate. Duplicate peak locations are slightly jittered for visualisation. Vertical and horizontal lines mark reward consumption at 0 cm on proximal and distal trials, respectively. The yellow dashed rectangle denotes the reward-alignment criterion: peak activity within *−*20 to +30 cm of reward on both proximal and distal trials. **F, G:** Same as **A, B** for mice in the second and third behavioural tercile, respectively. **H-I:** Activity of example place cells, **H**: two position-stable cells for a reward-reponsive mouse (J15), and **I**: two reward-aligned cells for a mouse in the reward-anticipatory tercile. Colourbars indicate the value of the calcium activity. On each reward-aligned plot (left hand side), *x* axis denote the position relative to the reward consumption, the green vertical line denotes the 0 origin of the reward consumption, the orange stars the reward location delivery with respect to consumption. On each position-aligned plot (right hand sides), each row is a trial (*y* axis), and the *x* axis is the position along the track, yellow vertical shaded area denotes the proximal reward consumption zone, orange shaded area the distal consumption zone. Pink stars denote the reward delivery location and purple stars the reward consumption location.

The unannounced relocation of reward beyond the familiar broad distribution provided a stronger test of whether strategy-dependent reward alignment would be maintained. Pooled and single-mouse maps before and after the switch in reward location indeed showed greater preservation of reward-aligned organisation towards the more spatially anticipatory end of the continuum (Fig. 3A–C and Supplementary Fig. S5). Using analogous criteria across the switch, the fraction of cells aligned near reward before and after relocation increased strongly with the anticipatory licking index (*β* = 4.22; exact mouse-level permutation, two-sided *p*_perm_ = 0.0057; Fig. 3D–G). Position stability was again not reliably associated with the anticipatory licking index in the primary continuous mouse-level analysis (exact mouse-label permutation test, two-sided *p*_perm_ = 0.1356; Supplementary Fig. S6D–G). We did not find any relationship between the anticipatory lick index and the percentage of cells which remap randomly (neither reward-aligned nor position stable; *β* = *−*0.37, *p*_perm_ = 0.84; Supplementary Fig. S6M). Moreover, reward alignment before relocation predicted reward alignment across it (*β* = 1.31; exact mouse-label permutation test, two-sided *p*_perm_ = 0.0009; Fig. 3H), while pre-switch position stability predicted position stability across the switch (*β* = 0.77; exact mouse-label permutation test, two-sided *p*_perm_ = 0.0203; Supplementary Fig. S6H).

**Fig. 3.**
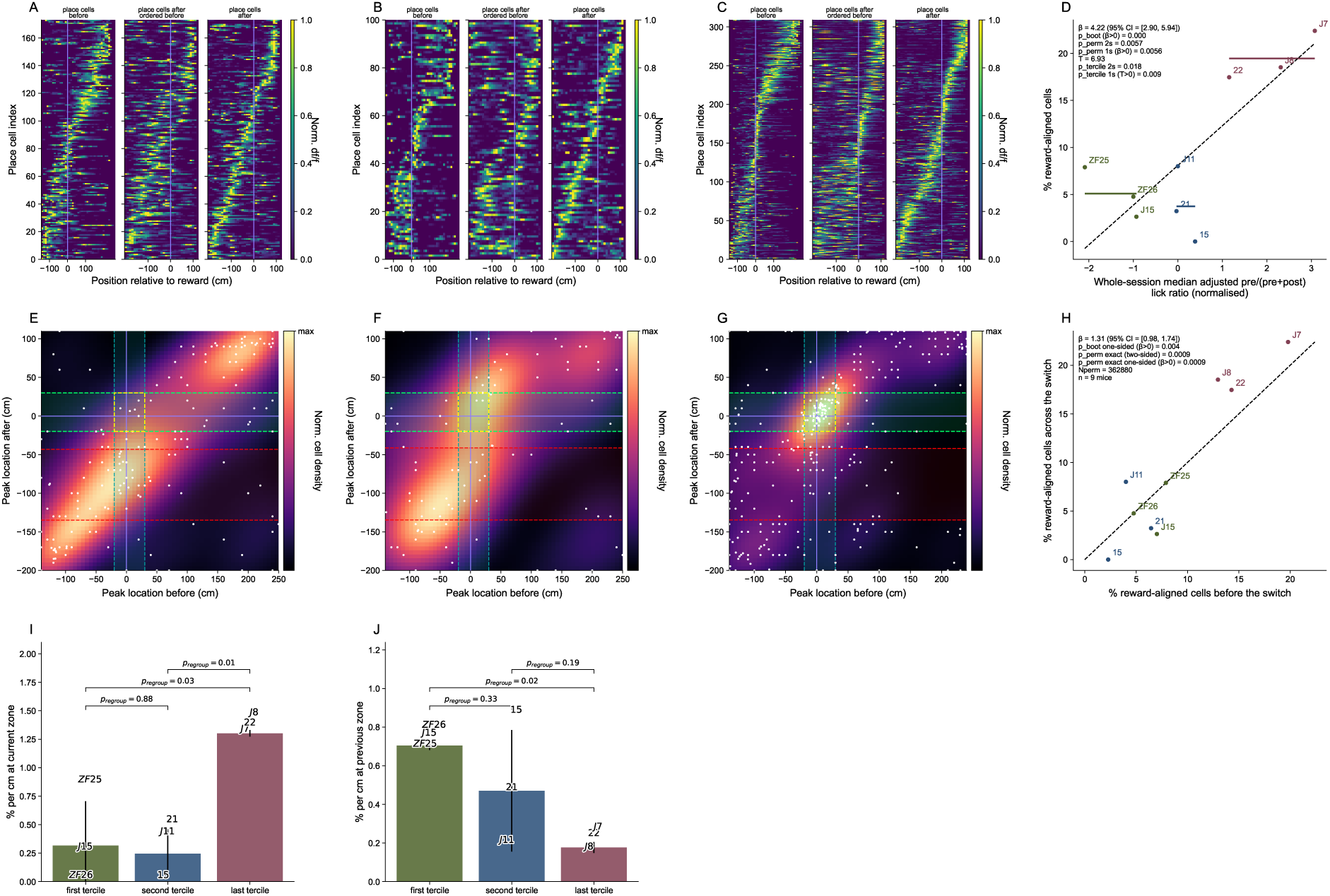
Reward-seeking strategy distinguishes persistent from flexible hippocampal reward coding across a reward-location switch. **A:** Normalised cross-validated reward-aligned place maps for the first behavioural tercile, for reward-responsive mice. Each row represents a cell that met the spatial-information criterion for a significant place cell both before and after the reward-location switch (see Methods). The left plot shows average activity on odd trials before the switch, ordered by the peak measured on even trials before the switch; the middle plot shows activity on odd trials after the switch while retaining the before-switch ordering; and the right plot shows activity on odd trials after the switch reordered by the peak measured on even trials after the switch. Activity is aligned to reward consumption and normalised to each cell’s peak. The vertical line marks reward consumption at 0 cm. **B, C:** Same as **A**, respectively, for mice in the second and third behavioural terciles, respectively. **D:** Across-mouse relationship between the normalised adjusted pre/(pre+post) lick ratio (*x* axis) and the proportion of cells that remained reward-aligned across the reward-location switch (*y* axis). Cells included in this analysis have peak locations in the yellow squares in **E, F, G** and in supplementary Figure S5D. Each point represents one mouse and is labelled by mouse identity; colours indicate lick-ratio terciles. Horizontal bars show the mean proportions of reward-aligned cells within each tercile. The dashed line shows the weighted linear-regression fit across mice (*β* = 4.22, 95% bootstrap CI [2.90, 5.94]; one-sided bootstrap test for *β >* 0, *p*_boot_ = 3.33 *×* 10*^−^*^4^; exact mouse-label permutation test: two-sided *p*_perm_ = 0.0057, one-sided *p*_perm_ = 0.0056). The ordered-tercile statistic was *T* = 6.93 (exact ordered-tercile permutation test: two-sided *p*_tercile_ = 0.0179, one-sided *p*_tercile_ = 0.0089; 1,680 ordered 3/3/3 assignments; *n* = 9 mice). **E:** Reward-aligned remapping across the reward-location switch for the same cells and mice as in **A**. The *x* and *y* coordinates show each cell’s peak reward-aligned activity location before and after the switch, respectively. Each white point represents one place cell, and the magma heat map shows a normalised Gaussian kernel-density estimate. Duplicate peak locations are jittered for visualisation. Vertical and horizontal blue lines mark reward consumption at 0 cm before and after the switch. The yellow dashed rectangle denotes the reward-aligned criterion: peak activity within *−*20 to +30 cm of reward both before and after the switch. Teal vertical dashed lines mark the previously reward aligned zone. Green horizontal dashed lines the boundary of the current reward zone used for quantification on panel **I** and red horizontal dashed line mark the boundaries of the previous reward zone used for quantification in panel **J**. **F, G:** Same as **E**, respectively, for mice in the third behavioural tercile, for reward-anticipatory mice. **H:** Relationship between reward alignment before the switch (*x* axis) and reward alignment across the switch (*y* axis). The *x* axis shows, for each mouse, the percentage of pre-switch place cells whose reward-aligned peak was within the reward window (*−*20 to +30 cm from reward) on both proximal and distal reward trials. The *y* axis shows the percentage of cells whose reward-aligned peak was within the same reward window both before and after the switch. Points indicate individual mice and are coloured by behavioural tercile. The dashed identity line indicates the identity function. Reward alignment before the switch predicted alignment across the switch (*β* = 1.31, 95% bootstrap CI [0.98, 1.74]; one-sided bootstrap test for *β >* 0, *p*_boot_ = 0.004; exact mouse-label permutation test: two-sided *p*_perm_ = 0.0009, one-sided *p*_perm_ = 0.0009; *n* = 9 mice; 362,880 permutations). **I:** Density of previously reward-aligned cells whose after-switch reward-aligned peak fell within the current reward vicinity, shown for the first reward-responsive tercile (green), second tercile (blue), and third reward-anticipatory tercile (purple). Bars show means across mice, error bars show the across-mouse standard deviation, and mouse labels indicate individual values. Brackets show pairwise mean differences evaluated under exact regrouping of the nine mice into ordered groups of three (1,680 assignments; two-sided tests): first versus second tercile, *p*_regroup_ = 0.88; first versus third tercile, *p*_regroup_ = 0.03; and second versus third tercile, *p*_regroup_ = 0.01. **J:** Density of previously reward-aligned cells whose after-switch reward-aligned peak remained near the previous reward location. Plotting conventions and exact 3/3/3 regrouping tests are as in **I**: first versus second tercile, *p*_regroup_ = 0.33; first versus third tercile, *p*_regroup_ = 0.02; and second versus third tercile, *p*_regroup_ = 0.19.

Finally, we examined the locations of the post-switch peak activity of just those cells with a pre-switch peak within *−*20 to +30 cm of reward (Supplementary Fig. S6I–K). For the more reward-responsive mice, the post-switch peaks remained concentrated more strongly near the former reward region; for the more spatially anticipatory mice, the post-switch peaks were concentrated more strongly near the new reward locations. Density near the current reward was similar in the more reward-responsive and middle terciles (*p*_regroup_ = 0.88), but higher in the most spatially anticipatory tercile than in either group (*p*_regroup_ = 0.03 and *p*_regroup_ = 0.01, respectively; Fig. 3I). Conversely, density within the former reward zone was higher in the more reward-responsive than in the most spatially anticipatory tercile (exact 3/3/3 regrouping test, two-sided *p*_regroup_ = 0.02; Fig. 3J). The continuous analyses were directionally consistent, although the two-sided mouse-label permutation tests did not reach significance (current-reward density: *β* = 0.24, two-sided *p*_perm_ = 0.0672, prespecified one-sided *p*_perm_ = 0.0436; former-reward density: *β* = *−*0.12, two-sided *p*_perm_ = 0.0505, prespecified one-sided *p*_perm_ = 0.0240; Supplementary Fig. S6N,O). Density outside both reward-related regions did not differ detectably across terciles (two-sided *p*_regroup_ *≥* 0.0881; Supplementary Fig. S6L), arguing against a general difference in remapping. Representative cells activity illustrate persistence and updating at the two ends of the continuum (Supplementary Fig. S7). Thus, profiles for the more reward-responsive mice were associated with greater persistence near the former reward, whereas stronger spatial anticipation was associated with greater updating towards the current reward.

These analyses reveal that spontaneous differences in reward-seeking strategy evoked by the same environment predicted the prevalence of reward-aligned neurons in CA1, along with the lability of alignment when the reward moved location unexpectedly, while substantial position coding was retained across animals. Thus, individual behavioural strategy provides a measurable axis along which hippocampal maps differ in the anchoring they treat as stable.

Our results complement work showing that behavioural engagement gates the reliability of hippocampal place codes [11]. Those authors distinguished engaged from disengaged trials using reward-zone lick selectivity and overall licking, and found that CA1 spatial coding degraded when mice disengaged despite traversing the same virtual environment. Our behavioural axis is different: all mice in our cohort showed spatially modulated pre-reward licking, indicating that they used spatial information and were engaged in the task, but they differed in the strength of the anticipatory component within an otherwise reward-localised behavioural profile. Thus, whereas engagement may determine whether CA1 maintains a reliable spatial code, variation from more reward-responsive to more spatially anticipatory reward seeking predicted the prevalence and updating of reward-aligned CA1 coding. This distinction extends work on hippocampal reward and goal coding [6, 7, 9, 10, 12] by showing that reward-aligned coding is shaped by how individual animals use those statistics to predict reward.

These strategy-dependent maps may reflect different ways in which hippocampal representations interact with attentional and action-selection circuits. One possibility is that, in more reward-responsive mice, hippocampal activity provides a spatial or contextual scaffold that identifies where reward-relevant cues should be sampled, while local outcome- or cue-triggered policies govern the final licking response. By contrast, anticipatory mice may rely more strongly on hippocampal outputs to ventral striatum, to convert spatial reward predictions into appetitive actions [13]. This interpretation is consistent with evidence that attention to spatial context stabilises hippocampal maps [14], and with classical and modern views that hippocampal and striatal systems support different but interacting navigation strategies [15–17]. Because the one-dimensional virtual track constrained overt behaviour largely to forward locomotion and licking, future work should manipulate strategy within animals and test the same relationship in tasks permitting richer navigation policies. Such experiments will be needed to determine whether behavioural strategy drives CA1 organisation, reflects it, or arises with it from a shared estimate of reward statistics. Regardless of directionality, individual reward-seeking strategy provides a behavioural window onto how CA1 organises variable and changing experience.

More broadly, our findings suggest that the nature and lability of hippocampal maps reflect the interpretation of space revealed by the structure of behaviour in that space. Behavioural strategy may therefore provide a readout of the internal model organising CA1 activity, linking reward expectation to the stability and updating of hippocampal representations.

## Online Methods

### Dataset and relationship to previous study

The virtual-reality task was developed for a related uncertainty study [10] which included two mice. The present manuscript analyses nine mice recorded in the same broad-zone switch condition, comprising those two initial mice and seven additional animals, and asks whether within-condition variation in reward-seeking strategy predicts CA1 organisation.

### Animals and surgery

All experiments were approved by, and conducted in accordance with, the Northwestern University Animal Care and Use Committee. Nine male P56–P63 C57BL/6J mice (Jackson Laboratory, stock no. 000664) were used. To express the calcium indicator jGCaMP8m, mice were injected with AAV1-syn-FLEX-jGCaMP8m-WPRE [18] into right dorsal CA1 (1.8 mm lateral, 2.3 mm caudal to Bregma, 1.25 mm below the dura). After injection, mice recovered with ad libitum water for 1–2 d and were then water restricted to 0.8–1.2 ml per day. Body weight was monitored and maintained at 75–80% of the original weight.

After 3–5 d of water restriction, hippocampal cannula implantation was performed above the injection site to allow optical access to dorsal CA1, as previously described [19]. Briefly, cortex above the dorsal hippocampus was aspirated until the white matter of the external capsule was exposed. Phosphate-buffered saline was applied repeatedly until bleeding stopped, a small drop of Kwik-Sil was applied to the tissue surface, and the cannula was inserted. A head plate and ring were cemented to the skull using Meta-bond. Analgesic and anesthetic procedures followed the approved animal protocol. Mice recovered for 5–7 d before behavioural training.

### Virtual reality task

Mice were habituated for one 45 min session in the head-fixed virtual reality setup [20] with the screen off, during which water rewards were delivered randomly to familiarize animals with the lick port. Beginning with the second session, the virtual reality screens were turned on and mice were trained in one visual environment to perform the task as described in the next paragraph. Training sessions lasted 45 min to 1 h depending on the number of laps run. Mice were considered trained when they ran at least 1–2 laps per minute, showed anticipatory licking before reward on at least 50% of laps, and exhibited stable behaviour across three consecutive sessions, measured by the average correlation coefficient of velocity and licking patterns across laps. All mice reached this criterion after 8–10 training sessions.

Head-fixed mice (*N* = 9) on a wheel ran in 1d virtual reality (VR) environments in which water reward was initially delivered on each trial within a 1m reward zone once per traversal of a 3m long linear track (and could subsequently be consumed anywhere by licking). On every run, the reward location was randomly drawn uniformly from 10 potential locations evenly spaced within the 1m wide zone (Fig. 1D). Mice experienced one session per day (mean 88 runs *±*15 std) until their behaviour was stable. After training, mice experienced a switch session. Initial trials (*n*; on average *n* = 36.8 *±* 4.5 std) in the session had the same location contingencies as those experienced in previous sessions. On trial *n* +1, without prior notice, the reward delivery location switched to one of two positions at the edges of a more remote 10cm zone. Behavioural variables, including licking, linear track position, velocity and reward delivery, were synchronized with two-photon imaging.

### Two-photon calcium imaging

Two-photon calcium imaging of dorsal CA1 neurons was performed using a custom-built movable-objective microscope with a 40*×*/0.8 NA water-immersion objective (LUMPlanFL N 40*×*/0.8 W, Olympus), as described previously [19, 20]. Scanning was controlled with ScanImage 5.1 (Vidrio Technologies). Average laser power after the objective was approximately 60–100 mW. Time-series movies of 12,000–24,000 frames were acquired at 512 *×* 256 pixels and approximately 30 Hz. A Digidata 1440A acquisition system with Clampex 10.3 (Molecular Devices) recorded behavioural variables at 1 kHz and synchronized them with imaging frame timing. The imaging field was kept constant within a session; across consecutive imaging sessions, imaging fields were not identical, although they could overlap.

### Image processing and calcium-event extraction

Two-photon movies were processed in Suite2p [21] for rigid and non-rigid motion correction and extraction of putative cell regions of interest. ROI fluorescence traces were exported to MATLAB for extraction of significant calcium transients [19]. Neuropil contamination was subtracted from each raw fluorescence trace after multiplication by 0.7. Slow fluorescence changes were removed by subtracting the eighth percentile of the fluorescence distribution in a 20 s window around each time point. Significant transients were identified by analyzing the ratio of positive- to negative-going transients across amplitudes and durations, yielding an estimated false-positive rate below 1%. Significant transients were retained and all other values were set to zero. These traces were used as changes in fluorescence for subsequent analyses.

### Place-cell identification

Fluorescence tuning maps were generated by binning position across the track into 60 spatial bins and calculating, for each bin, the mean event-filtered calcium fluorescence signal during periods in which the animal moved at least 0.1 cm s*^−^*^1^. Spatial information, *I*, was calculated in bits per event-filtered fluorescence [22]:

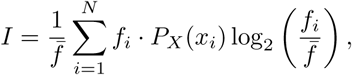

where *f̅* is the mean event-filtered fluorescence signal, *N* is the number of spatial bins, *f_i_* is the mean event-filtered fluorescence signal in the *i*th spatial bin, and *P_X_* (*x_i_*) is the occupancy proportion of that bin. A null distribution was generated by circularly shuffling each fluorescence trace with a minimum shift of 15 s and recalculating the tuning map 1,000 times. A cell was classified as a significant place cell if its spatial information exceeded 99% of values from the shuffled null distribution and was at least 0.5 bits per event-filtered fluorescence signal.

### Behavioural and place-cell analyses in position and reward-consumption-aligned coordinates

Lick-rate and velocity traces were downsampled at 30 Hz and averaged across 10 cm position bins weighted by occupancy. Session averages before the switch were computed separately for proximal, middle and distal reward trials and normalized to the maximum value of the full-session average (Fig. 1G and Supplementary Fig. S2).

Reward-consumption position was defined as the position of the first lick after reward delivery. The reward zone was defined from the most proximal to the most distal reward consumption position. Trials without reward consumption, or in which reward consumption occurred more than or exactly 12.38 cm after reward delivery, were excluded from reward-aligned behavioural and neural analyses (Supplementary Fig. S1). In total, 35 (4.0%) of 878 trials were excluded; the distribution of consumption delays and the excluded trials are shown in Supplementary Fig. S1E.

Proximal, middle and distal reward trials were defined by dividing the reward consumption zone into three bins of identical length: the maximum and minimum of reward consumption locations were considered as reward zone and this distance was divided by 3. Trials in which reward was consumed in the first, second or third bin were labeled proximal, middle or distal, respectively. This lead to an average of 12.78 *±* 3.60 proximal, 12.22 *±* 3.23 middle, and 11.56 *±* 2.74 distal trials per mouse respectively.

Lick-rate and velocity visualisations in Fig. 1E,G and Supplementary Fig. S2A–C,E–G were aligned to reward consumption. Reward-aligned place-cell visualisations and analyses in Figs. 2 and 3 and Supplementary Figs. S3, S5 and S6 were likewise computed relative to reward consumption. By contrast, the adjusted pre/(pre+post) licking index in Fig. 1F and Supplementary Fig. S1A–C was defined relative to reward delivery, because it quantified licking before versus after reward became available.

Teleportation periods were excluded from all analyses. Behavioural analyses also excluded data points with velocity below 0.1 cm s*^−^*^1^; place-cell analyses excluded periods with velocity below 1 cm s*^−^*^1^. Behavioural variables and calcium activity were averaged over 10 cm spatial bins, weighted by time spent in each bin. Place maps were cross-validated by averaging activity on odd trials and ordering cells according to peak activity on even trials. For switch sessions, maps before and after the switch were computed separately and normalised by each cell’s maximum activity.

Peak activity was defined as the 10 cm bin with maximum average activity. Stability was evaluated across proximal and distal reward trials before the switch (Fig. 2 and Supplementary Figs. S3 and S4), or across trials before and after the reward-location switch (Fig. 3 and Supplementary Figs. S5 and S6). Reward-consumption-aligned coordinates were computed relative to reward consumption on each trial using 10 cm bins.

### Anticipatory lick ratio or Pre/post reward licking index

Anticipatory licking was quantified by an adjusted pre/(pre+post) lick ratio (Fig. 1F and Supplementary Fig. S1A–C). For each phase and mouse, reward-delivery-aligned lick-rate traces were used to estimate the distance from reward delivery to the peak of the mean post-reward lick response, with a minimum distance of 20 cm. Pre- and post-reward windows of 1.5 times this distance were then used to compute mean pre- and post-reward lick rates on each trial; the ratio was pre/(pre+post). Mouse-level values were medians across included trials.

### Cell-class definitions

For proximal-distal reward analyses, position-stable cells were place cells whose peak activity positions on proximal and distal reward trials differed by at most 30 cm (Supplementary Figs. S3E, limegreen dashed lines, and S4E–G). Reward-aligned cells were defined as cells with reward-consumption-aligned peaks within *−*20 to +30 cm of reward consumption on both proximal and distal trials, after excluding position-stable cells (Fig. 2E–G and Supplementary Fig. S3D, area shown with limegreen dashed rectangle).

Position-stable cells across the switch were defined as place cells whose before- and after-switch peak locations differed by less than 30 cm in absolute-position coordinates (Supplementary Figs. S5E and S6E–G).

Reward-aligned cells were defined as cells with reward-consumption-aligned peaks within *−*20 to +30 cm of reward consumption both before and after the switch, after excluding position-stable cells (Fig. 3E–G and Supplementary Fig. S5D).

### Statistics

Statistical analyses were performed using custom Python code with `numpy`, `scipy` and `statsmodels`. Unless otherwise stated, the mouse was the unit of inference for analyses relating behavioural strategy to neural coding. For each mouse, anticipatory licking was summarized as the median adjusted pre/(pre+post) lick ratio across included trials. Continuous mouse-level relationships between licking and neural measures were quantified by linear regression (Fig. 2D, Fig. 3D,H and Supplementary Figs. S4D and S6D,H,M–O). For the main cell-class analyses, slopes were estimated by weighted least squares, with mouse weights defined as the product of the number of included pre-switch behavioural trials and the number of analysed cells; equal-weight and cell-count-only fits were used as sensitivity analyses where indicated. Slope uncertainty was estimated from 3,000 bootstrap resamples of mice.

Permutation-based *p* values were used for mouse-level slope analyses (Fig. 2D, Fig. 3D,H and Supplementary Figs. S4D and S6D,H,M–O). For the primary nine-mouse weighted regressions, exact permutation tests enumerated all 9! = 362,880 possible assignments of the nine mouse-level neural outcomes shown on the y-axis across the nine fixed behavioural values and regression weights. Tercile analyses assigned the nine mice to low, middle and high anticipatory-licking groups of three mice each and used exact ordered-tercile permutation tests over all 1,680 possible 3/3/3 assignments (Fig. 2D, Fig. 3D and Supplementary Figs. S4D and S6D,M–O). Full details of the weighting, bootstrap intervals, permutation *p* values, tercile tests and spatial-density comparisons are provided in Supplementary Methods.

Spatial-density comparisons of post-switch peak locations used mouse-level exact paired sign-flip tests for within-tercile comparisons and exact regrouping tests for across-tercile comparisons (Fig. 3I,J and Supplementary Fig. S6L,N,O). Proportions were compared using proportion z-tests only for pooled cell-count comparisons; one-sided tests were used only for prespecified directional comparisons. Distributions were compared using Kolmogorov–Smirnov tests. Gaussian kernel density estimates were used only for visualization of two-dimensional peak-location distributions (Fig. 2, Fig. 3, Supplementary Fig. S3 and Supplementary Fig. S5). Unless otherwise stated, statistical tests were two-sided and *p* values were not adjusted for multiple comparisons. Subscripts identify the resampling procedure: *p*_boot_ denotes bootstrap tail probabilities, *p*_perm_ denotes mouse-label permutation tests of continuous regression slopes, *p*_tercile_ denotes ordered-tercile permutation tests, *p*_regroup_ denotes pairwise contrasts evaluated under exact 3/3/3 mouse regrouping, and *p*_shuffle_ denotes within-mouse shuffle tests.

## Data availability

Processed data supporting the figures will be made available upon publication and are available from the corresponding authors upon reasonable request.

## Code availability

Analysis code will be made available upon publication and is available from the corresponding authors upon reasonable request.

## Acknowledgements

We are very grateful to Daniel Dombeck for providing access to the dataset, to him and Jack Mellor for invaluable discussions, conceptual development, scientific input throughout the project, together with task design and implementation, and to Feng Xuan and Zehua Chen for behavioural and imaging data acquisition.

This work was supported by the Simons Foundation, the Wellcome Trust, the Max Planck Society, and the Alexander von Humboldt Foundation.

## Author contributions

C.T. developed the analytical framework, performed all analyses, interpreted the results, prepared the figures and wrote the manuscript. P.D. contributed to the conceptual development of the study, interpretation of the results, supervision, and manuscript writing and editing.

---

**Fig. S1.**
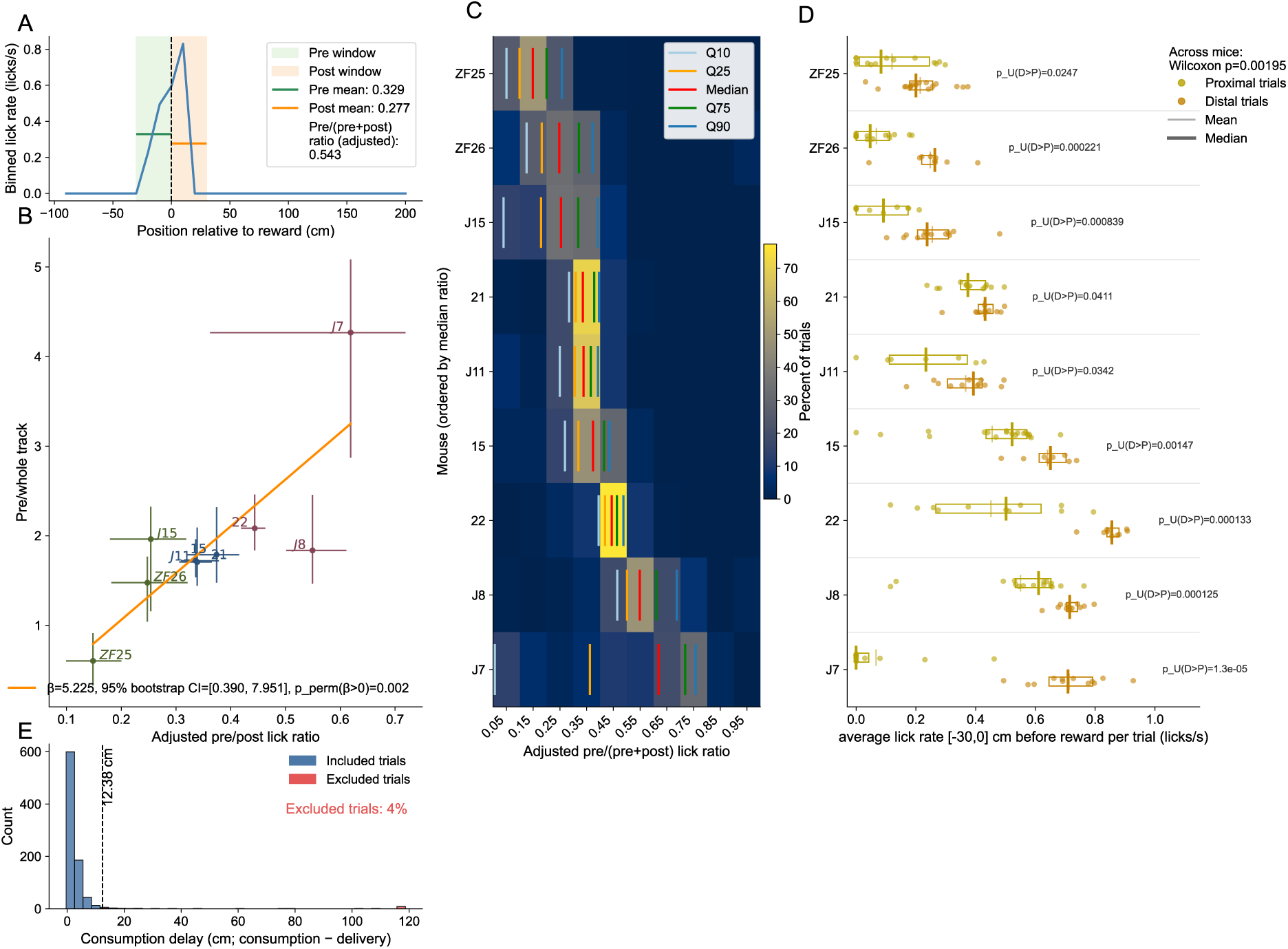
Quantification and validation of anticipatory licking, relationship with another selective licking metric, and trial inclusion criteria. **A:** Example trial illustrating the computation of the adjusted anticipatory licking metric. The blue trace is the lick rate for this trial. For each animal and each phase (before and after the switch separately), we first estimate a reward-response distance from the position of the peak lick rate after reward delivery. The post-reward window was then defined from reward delivery to 1.5*×* this reward-response distance after reward delivery (orange shaded region), and the pre-reward window was defined as the matched window of identical length immediately preceding reward delivery (green shaded region). The adjusted pre/(pre+post) lick ratio was computed as the mean lick rate in the pre-reward window divided by the sum of the mean lick rates in the pre- and postreward windows. The pre/whole-track metric used in B was computed as the mean lick rate in the pre-reward window divided by the sum of the mean lick rate in the pre-reward window and the mean lick rate over the remainder of the track. **B:** The adjusted anticipatory licking ratio agrees with a whole-track-normalised anticipatory lick-selectivity measure used in recent VR navigation studies [9]. Each point corresponds to one animal (*n* = 9 mice), plotted as the session median adjusted pre/(pre+post) lick ratio (*x* axis) against the session median pre/whole-track licking ratio (*y* axis). Horizontal and vertical whiskers denote the interquartile range across trials. The orange line shows the linear fit (*β* = 5.225, 95% bootstrap CI [0.390, 7.951]; one-sided exact mouse-label permutation test, *p*_perm_ = 0.0022). **C:** Distribution of the adjusted pre/(pre+post) lick ratio across all trials for each animal over the full session, including trials before and after the switch. Animals are ordered by their median adjusted pre/(pre+post) lick ratio. Heat-map values indicate the percentage of trials falling in each ratio bin (*x* axis). Coloured vertical lines denote the 0.1, 0.25, 0.5, 0.75 and 0.9 quantiles for each animal. **D:** Spatial modulation of anticipatory licking before the switch in reward zone. For each animal, trials were separated into proximal and distal groups based on reward-consumption location, and the average lick rate in a reward-aligned pre-reward-delivery window of [*−*30, 0] cm was computed for each trial. Each dot represents one trial. Thin vertical lines indicate the mean, thick vertical lines indicate the median, and rectangles indicate the interquartile range. Animals are shown in the same order as in **C**. *P −*values shown for each animal are one-sided Mann–Whitney *U* tests comparing distal versus proximal trials, testing whether distal trials had higher pre-reward lick rates than proximal trials. Across mice, the population-level effect was assessed by first computing the distal-proximal mean difference for each mouse and then applying a one-sided Wilcoxon signed-rank test (*p* = 0.00195). **E:** Distribution of reward-consumption delays across all trials of the nine animals, pooling trials before and after the switch. Consumption delay was calculated as the reward-consumption position minus the reward-delivery position. The stacked histogram (40 bins) shows included trials in blue (delay *<* 12.38 cm) and threshold-excluded trials in red (delay *≥* 12.38 cm); the dashed black line indicates the 12.38 cm exclusion threshold. Overall, 35 of 878 trials were excluded (3.99%): 28 because their consumption delay reached or exceeded the threshold and seven because a valid reward-delivery or consumption position was unavailable. These seven trials contribute to the reported exclusion percentage but cannot be displayed in the histogram because their consumption delay could not be calculated.

**Fig. S2.**
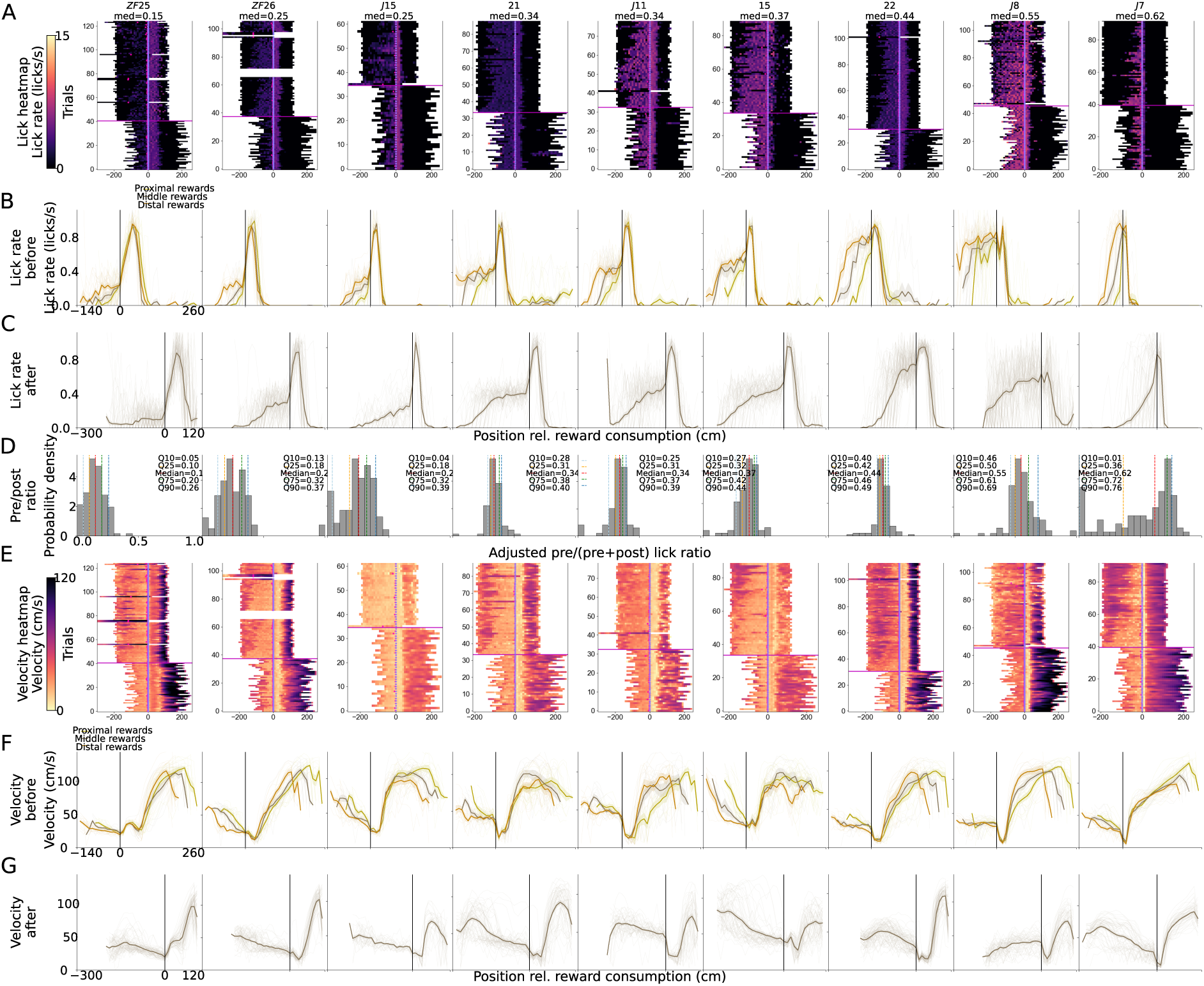
Single-animal licking profiles reveal behavioural heterogeneity. Each column corresponds to one animal, ordered from left to right by increasing median adjusted pre/(pre+post) lick ratio computed across the session, thereby arranging animals along a continuum from predominantly post-reward-responsive to relatively more anticipatory licking. Panels A–C and E–G are aligned to reward consumption, whereas the adjusted pre/(pre+post) index in D was computed relative to reward delivery as described in Supplementary Fig. S1A and Methods. Only trials satisfying the reward-consumption inclusion criterion were shown (Supplementary Fig. S1E and Methods). **A:** Reward-consumption-aligned lick-rate heatmaps for individual animals. Each horizontal row is one trial, and the magenta horizontal line marks the switch in reward contingency. The *x* axis shows position relative to reward consumption, such that 0 cm corresponds to the first lick after reward delivery. The vertical line marks reward consumption at 0 cm. Colour indicates lick rate (licks s*^−^*^1^), with the scale shown at the left. Magenta stars indicate reward-delivery positions relative to reward consumption. **B:** Mean lick-rate traces before the switch, aligned to reward consumption. Trials are grouped by reward-consumption location into proximal, middle and distal groups. Thin lines show individual trials, thick lines show the mean, and the vertical black line marks reward consumption at 0 cm. **C:** Mean lick-rate traces after the switch, aligned to reward consumption and plotted as in B. Thin lines show individual trials, thick lines show the mean, and the vertical black line marks reward consumption at 0 cm. **D:** Distribution of the adjusted pre/(pre+post) lick ratio for all included trials from the session shown in that column. Grey bars with black outlines show probability density (*y* axis). Vertical dashed lines mark the 10th (light blue), 25th (orange), 50th (red), 75th (green), and 90th (dark blue) quantiles; values are listed in the legend of each subplot. **E:** Reward-consumption-aligned velocity heatmaps for individual animals, arranged as in A. The vertical line marks reward consumption at 0 cm. Each horizontal row is one trial, the magenta horizontal line marks the switch in reward contingency, and magenta stars indicate reward-delivery positions relative to reward consumption. Colour indicates velocity (cm s*^−^*^1^), with the scale shown at the left. **F:** Mean velocity traces before the switch, aligned to reward consumption and plotted as in B. Yellow, grey-brown, and orange traces denote proximal, middle, and distal reward-location trials averages. Thin lines show individual trials, thick lines show the mean, and the vertical black line marks reward consumption at 0 cm. **G:** Mean velocity traces aligned to reward consumption after the switch, plotted as in F. Thin lines show individual trials, thick lines show the mean, and the vertical black line marks reward consumption at 0 cm.

**Fig. S3.**
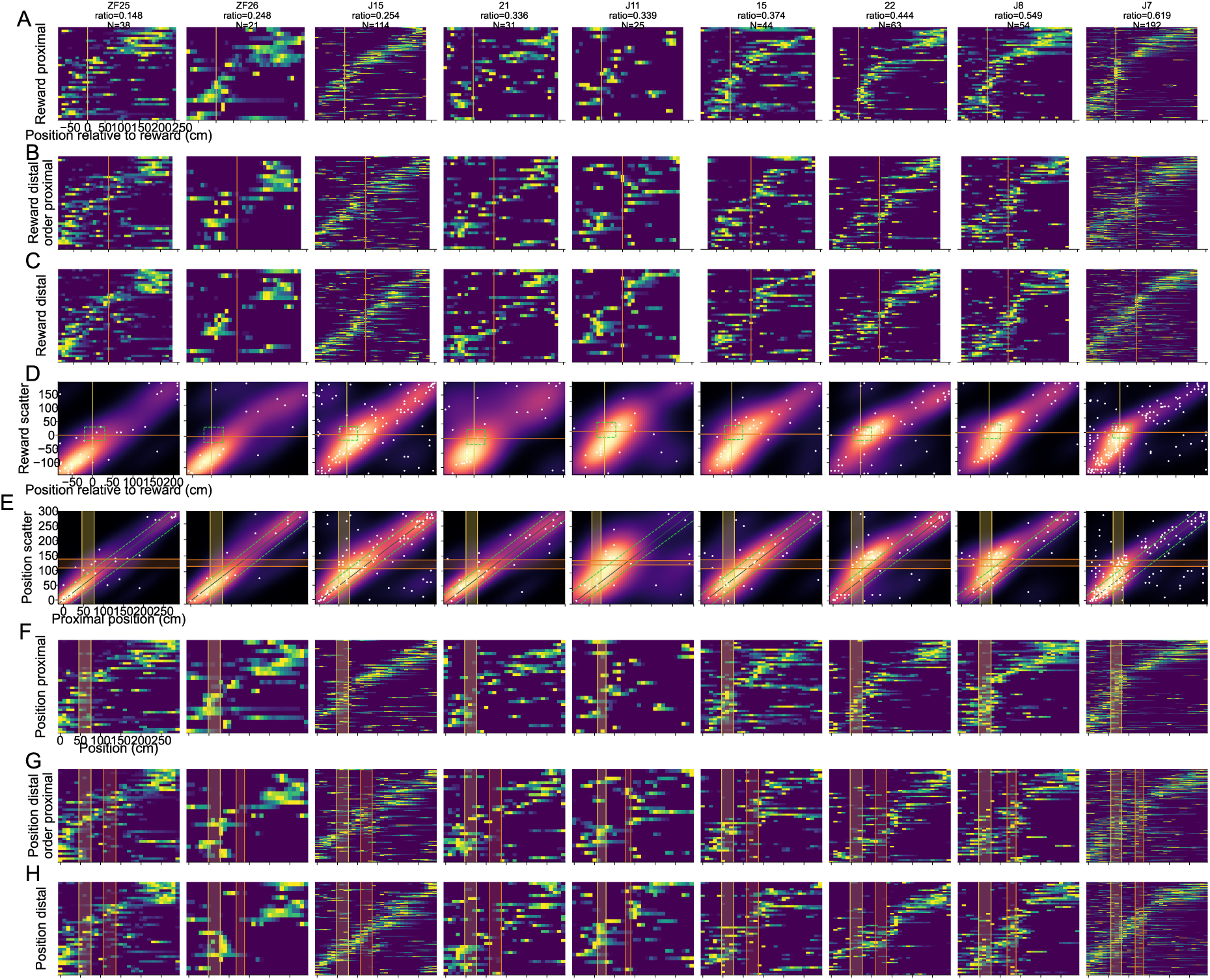
Single-animal reward- and position-frame stability across proximal and distal reward trials before the switch. Column titles show animal identity, median adjusted pre-to-post lick ratio, and the number of place cells included. Animals are ordered by their median adjusted pre-to-post lick ratio across all trials. Place cells were defined as cells passing the place-cell criterion before the switch. All panels in this figure use only trials before the switch, split into proximal and distal reward-location groups. Heatmaps show row-normalised mean activity (normalised Δ*F/F*) for each cell in 10 cm spatial bins. Heatmap activity is averaged over odd trials and cells are ordered by the peak location in the even-trial half of the corresponding trial group, except where explicitly displayed in proximal-trial order. **A–C:** Reward-consumption aligned maps. The *x* axis is position relative to reward consumption location, with 0 cm marking reward consumption. **A:** Normalised activity of shared place cells on proximal trials, ordered by each cell’s even-trial proximal peak in reward coordinates. **B:** Normalised activity of the same shared place cells on distal trials, displayed in the proximal-trial order to reveal whether the proximal reward-centred organisation is preserved or reorganised on distal trials. **C:** Normalised distal-trial activity reordered by each cell’s even-trial distal peak in reward coordinates. **D:** Reward-consumption aligned proximal-versus-distal remapping scatter plot. Each white point is one place cell, with *x* and *y* coordinates corresponding to the peak average activity location on proximal and distal trials, respectively. The magma heatmap shows the Gaussian kernel-density estimate of cell density; point jitter is applied only for visualisation of overlapping 10-cm binned peak locations. Vertical and horizontal reference lines mark the reward location (0 cm) for proximal and distal trials. The limegreen dashed square denotes the reward-aligned window used for Fig.2D analyses (*−*20 to +30 cm relative to reward on both axes). **E:** Position-reference-frame proximal-versus-distal remapping scatter plot for the same cells. The *x* and *y* axes show peak activity location in absolute track coordinates on proximal and distal trials, respectively. White points and magma shading are as in **D**. Vertical yellow shading and lines mark the proximal reward zone, orange horizontal shading and lines mark the distal reward zone, and limegreen dashed diagonals indicate *±*30 cm around the identity line, corresponding to the position-stability criterion quantified in Supplementary Fig. S4D. **F–H:** Position-reference-frame maps. The *x* axis is absolute position on the track. Yellow and orange shading and lines indicate respectively the proximal and distal reward zones. **F:** Normalised proximal-trial activity ordered by each cell’s even-trial proximal peak in position coordinates. **G:** Normalised distal-trial activity displayed in the proximal-trial order. **H:** Normalised distal-trial activity reordered by each cell’s even-trial distal peak in position coordinates.

**Fig. S4.**
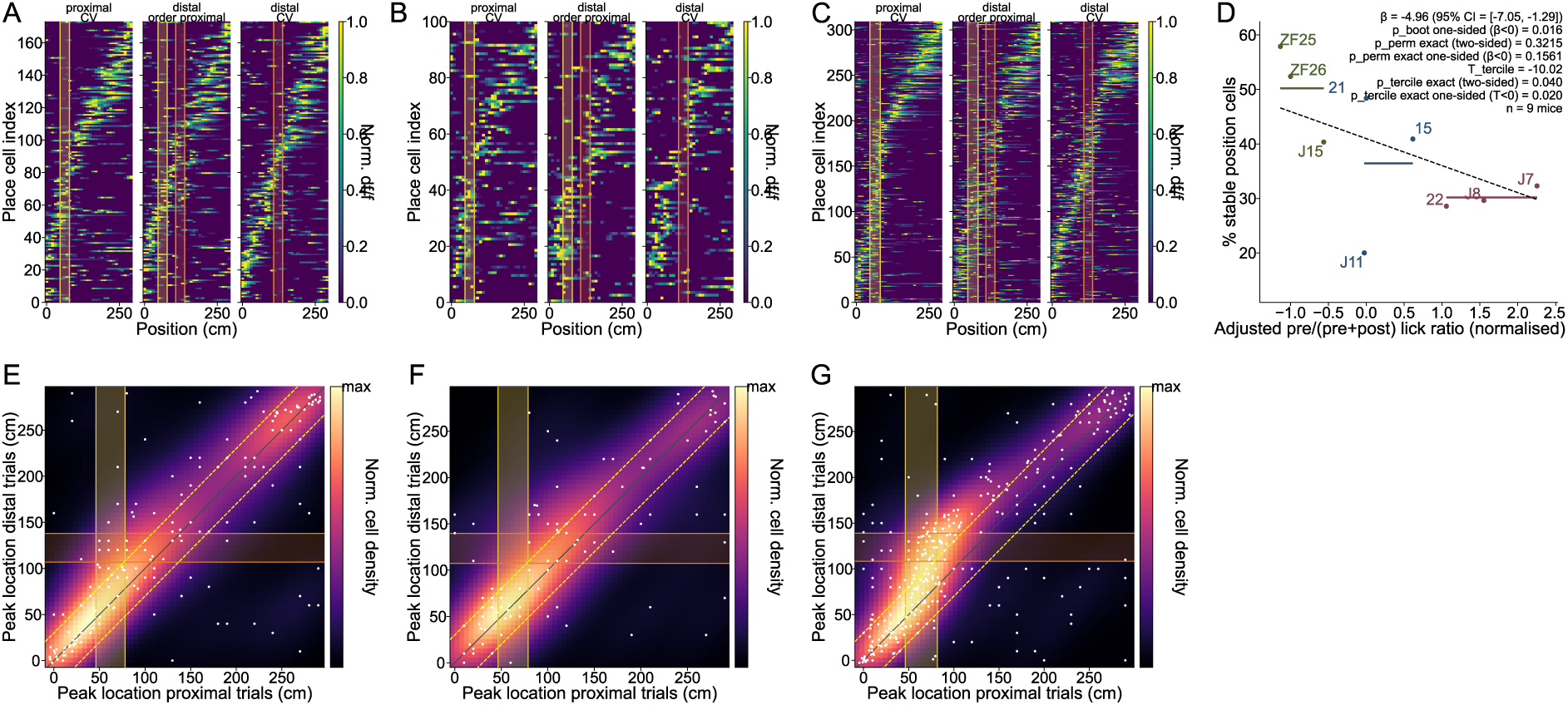
Position stability across variable reward locations. **A:** Normalised cross-validated position-aligned place maps for the first behavioural tercile, pooled across reward-responsive mice. Analyses use trials before the reward-location switch, divided according to reward-consumption location into proximal and distal reward trials. Each row represents a cell that met the spatial-information criterion for a significant place cell before the switch (see Methods). The left plot shows average activity on odd proximal trials, ordered by the peak activity measured on even proximal trials; the middle plot shows activity on odd distal trials while retaining the proximal-trial ordering; and the right plot shows activity on odd distal trials reordered by the peak measured on even distal trials. Activity is plotted in absolute position coordinates and normalised to each cell’s peak. Yellow and orange lines and shaded regions indicate the proximal and distal reward-consumption zones, respectively. **B, C:** Same as **A** for mice in the second and third behavioural terciles, respectively. **D:** Across-mouse relationship between the normalised adjusted pre/(pre+post) lick ratio (*x* axis) and the proportion of position-stable cells across proximal and distal reward trials before the switch (*y* axis). Position-stable cells were defined using the 30-cm proximal–distal peak-displacement criterion illustrated in **E–G**. Each point represents one mouse and is labelled by mouse identity; colours indicate lick-ratio terciles. Horizontal bars show the mean proportion of position-stable cells within each tercile, and the dashed line shows the weighted linear-regression fit across mice. The fitted weighted-regression slope was negative (*β* = *−*4.96, 95% bootstrap CI [*−*7.05*, −*1.29]), but the primary exact mouse-label permutation test did not support a reliable continuous association (exact mouse-label permutation test: two-sided *p*_perm_ = 0.3215, one-sided *p*_perm_ = 0.1561). The orderedtercile statistic was *T* = *−*10.02 (exact ordered-tercile permutation test: two-sided *p*_tercile_ = 0.040, one-sided *p*_tercile_ = 0.020; 1,680 ordered 3/3/3 assignments; *n* = 9 mice). **E:** Proximal-versus-distal position stability for the same cells and mice as in **A**. The *x* and *y* coordinates show each cell’s peak activity position on proximal and distal trials, respectively. Each white point represents one place cell, and the magma heat map shows a normalised Gaussian kernel-density estimate. The vertical yellow band indicates the proximal reward-consumption zone, whereas the horizontal orange band indicates the distal reward-consumption zone. The solid diagonal denotes identical peak positions on proximal and distal trials. Yellow dashed lines delimit the position-stability criterion: cells were classified as position-stable when their peak positions on proximal and distal trials differed by no more than 30 cm. **F, G:** Same as **E** for mice in the second and third behavioural terciles, respectively.

**Fig. S5.**
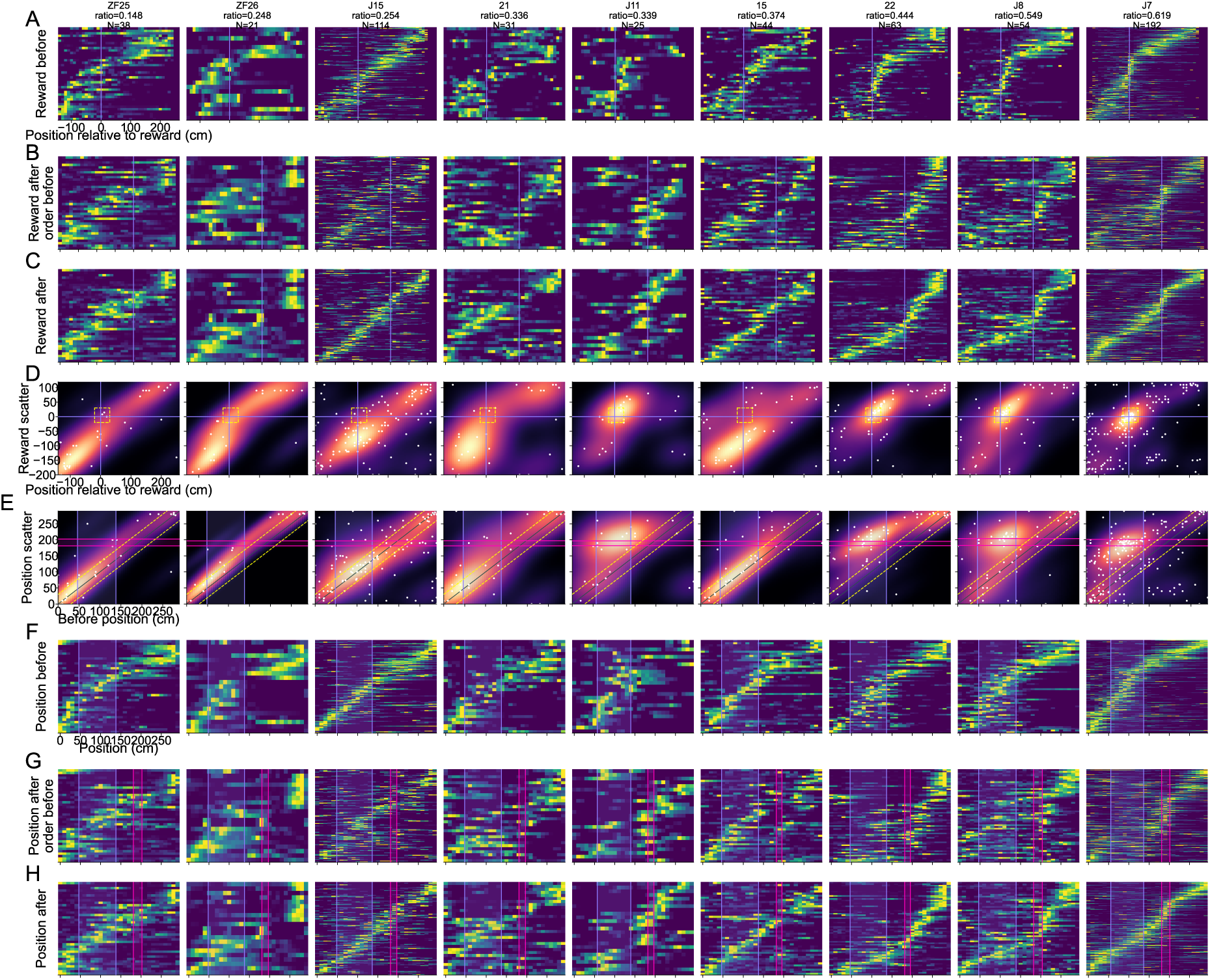
Single-animal hippocampal remapping profiles across reward relocation. Column titles show animal identity, median ratio, and the number of shared place cells included. Shared place cells were defined as cells passing the place-cell criterion in both the before- and after-switch epochs. Heatmaps show row-normalised mean activity (normalised Δ*F/F*) for each cell in 10-cm spatial bins. Activity is averaged and shown on odd trials for cells originally ordered based on their peak activity on even trials, except where explicitly ordered according to the before-switch map. **A–C:** Reward-consumption aligned maps. The *x* axis is the position relative to the reward consumption location, with 0 cm marking reward consumption. Purple vertical lines indicate reward alignment. **A:** Normalised before-switch activity of shared place cells, ordered by each cell’s even-trial before-switch peak in reward coordinates. **B:** Normalised after-switch activity of the same shared place cells, displayed in the before-switch order to reveal whether the pre-switch reward-centred organisation is preserved, displaced, or reorganised after reward relocation. Purple line denotes the post-switch reward aligned (0) coordinate. **C:** Normalised after-switch activity reordered by each cell’s even-trial after-switch peak in reward coordinates. **D:** Reward-consumption aligned remapping scatter plot. Each white point is one shared place cell, with *x* and *y* coordinates corresponding to the peak average activity location before and after the switch, respectively. The magma heatmap shows the Gaussian kernel-density estimate of cell density; point jitter is applied only for visualisation of overlapping 10-cm binned peak locations. Purple vertical and horizontal lines mark the reward location before and after the switch. The yellow dashed square denotes cells with peaks within the reward-centred window before and after the switch (*−*20 to +30 cm from reward). **E:** Position-reference-frame remapping scatter plot for the same cells. The *x* and *y* axes show all-trial peak activity location in absolute track coordinates before and after the switch, respectively. White points and magma shading are as in D. Purple vertical shading/lines mark the before-switch reward zone, magenta horizontal shading/lines mark the after-switch reward zone, and yellow dashed diagonals indicate *±*30 cm around the identity line, corresponding to position-stable peak locations. **F–H:** Position-reference-frame maps. The *x* axis is absolute position on the track. Purple shading/lines indicate the before-switch reward zone and magenta shading/lines indicate the after-switch reward zone. **F:** Normalised before-switch activity ordered by the even-trial before-switch peak in position coordinates. **G:** Normalised after-switch activity displayed in the before-switch order. **H:** Normalised after-switch activity reordered by the even-trial after-switch peak in position coordinates, showing the post-switch position map structure.

**Fig. S6.**
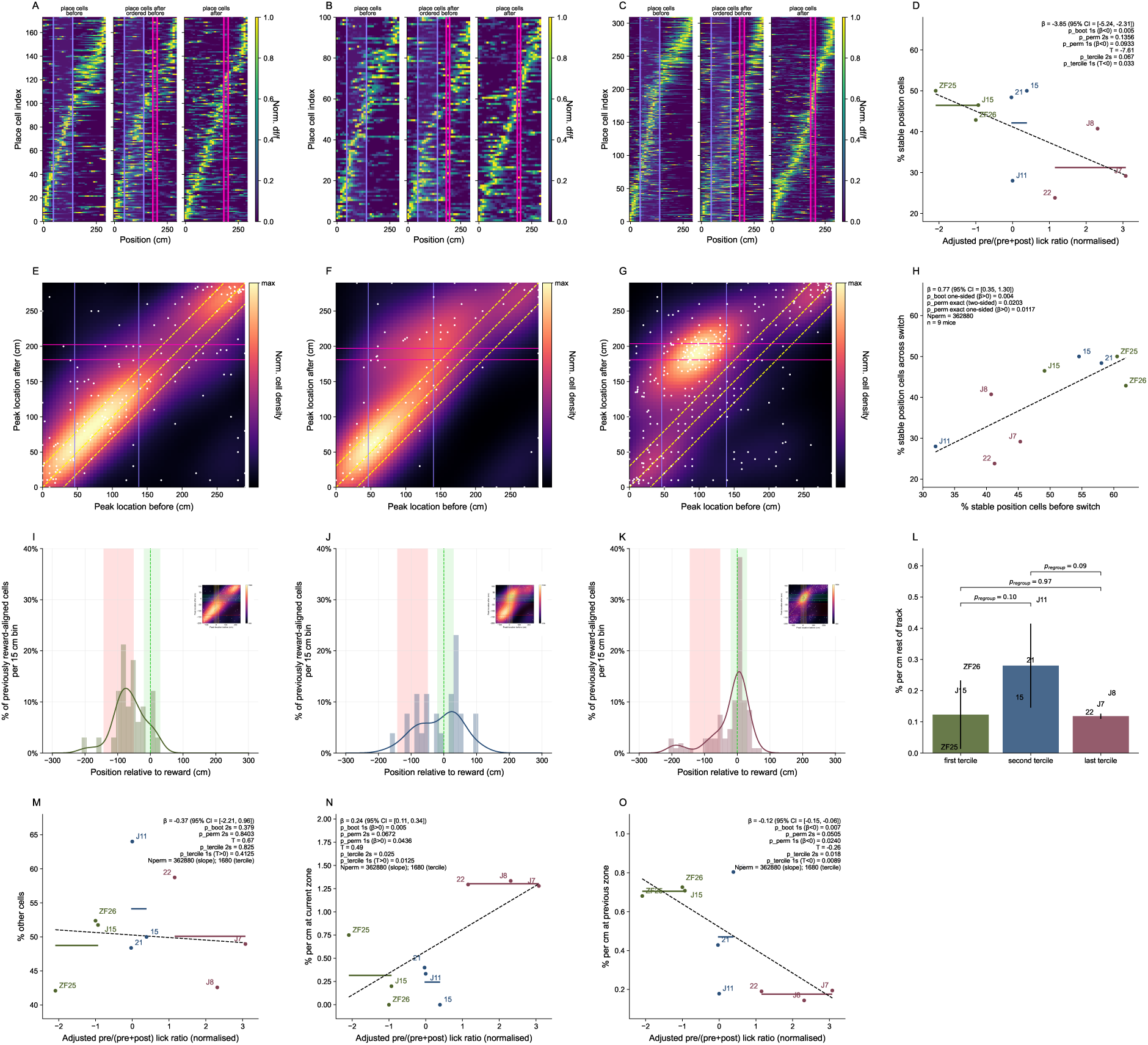
Reward-remapping profiles and position-reference-frame stability across behavioural terciles. **A–C:** Normalised cross-validated place maps. Each row is a cell that met the spatial-information criterion for being a significant place cell (see Methods) both before and after the reward-location switch. The left-hand plot shows the average activities of the cells on odd trials before the switch, ordered by their peak activity averaged on even trials before the switch; the middle plot shows the activity of the cells after the switch, ordered by their peak activity averaged on even trials before the switch; and the right-hand plot shows the average activity of the cells on odd trials after the switch, ordered by their peak activity averaged on even trials after the switch. Each cell’s activity is normalised to its peak. Vertical lines denote the reward zones: blue before the switch and pink after the reward-zone switch. **A:** The more reward-responsive tercile of mice; **B:** the middle tercile; and **C:** the most spatially reward-anticipatory tercile. **D:** Relationship between anticipatory licking and position stability across the reward switch. The *x* axis shows the normalised adjusted pre/(pre+post) lick ratio, and the *y* axis shows the across-switch position-stability measure illustrated in **E–G**. Points represent individual mice, are labelled by mouse identity and are coloured by lick-ratio tercile; horizontal bars show tercile means. The dashed line shows the weighted linear-regression fit. The fitted slope was negative (*β* = *−*3.85, 95% bootstrap CI [*−*5.24*, −*2.31], one-sided bootstrap test for *β <* 0, *p*_boot_ = 0.005), but the exact mouse-label permutation test did not support a reliable continuous association between anticipatory licking and position stability (exact mouse-label permutation test: two-sided *p*_perm_ = 0.1356, one-sided *p*_perm_ = 0.0933). The ordered-tercile statistic was *T* = *−*7.61 (exact ordered-tercile permutation test: two-sided *p*_tercile_ = 0.067, one-sided *p*_tercile_ = 0.033; *n* = 9 mice; 1,680 ordered 3/3/3 assignments). **E–G:** Scatter plots showing the positions of peak activity before (*x* axis) versus after the switch (*y* axis). Each white dot is a place cell and the heat map shows a Gaussian probability-density estimate (see Supplementary Material). Yellow dashed lines delineate the diagonal region used to quantify position-stable cells for the statistics in panel D. Scatter plots include jitter proportional to cell density, enhancing visualisation of overlapping data points. Vertical blue lines denote the reward zones before the switch, and pink horizontal lines denote the reward zone after the switch. **E:** The more reward-responsive tercile; **F:** the middle tercile; and **G:** the most spatially reward-anticipatory tercile. **H:** Relationship between position stability before and across the reward switch. Pre-switch position stability (*x* axis) was defined as the percentage of place cells whose peak locations in the position reference frame differed by no more than 30 cm between proximal- and distal-reward trials. Across-switch position stability (*y* axis) was defined as the percentage of cells whose position-frame peak locations differed by no more than 30 cm before versus after the switch. Points represent individual mice, are labelled by mouse identity and are coloured by lick-ratio tercile. The dashed line shows the linear-regression fit. Greater pre-switch position stability predicted greater across-switch position stability (*β* = 0.77, 95% bootstrap CI [0.35, 1.30]; one-sided bootstrap test for *β >* 0, *p*_boot_ = 0.004; exact mouse-label permutation test: two-sided *p*_perm_ = 0.0203, one-sided *p*_perm_ = 0.0117; *n* = 9 mice; 362,880 permutations). **I–K:** Distribution of post-switch reward-aligned peak activity locations among cells whose pre-switch reward-aligned peak lay within the reward vicinity (*−*20 to +30 cm relative to reward). Histograms show the percentage of selected cells per 15-cm bin for mice grouped according to their normalised adjusted pre/(pre+post) lick ratio: **I:** first tercile (reward-responsive; *n* = 33 cells); **J:** second tercile (*n* = 26 cells); and **K:** last tercile (reward-anticipatory; *n* = 107 cells). Curves show Gaussian kernel-density estimates of individual-cell peak locations. Red shading denotes the previous reward zone, green shading denotes the post-switch reward vicinity (*−*20 to +30 cm relative to the post-switch reward), and the green dashed line indicates the post-switch reward location. Insets show the corresponding pre-switch versus post-switch reward-aligned peak-location distributions; the horizontal shaded regions indicate the reward-zone criteria used to classify cells. **L:** Density of the selected, previously reward-aligned cells whose post-switch reward-aligned peak fell outside both the previous and current reward zones (“rest of track”). Density was calculated for each mouse as the percentage of selected cells divided by the length of the corresponding track region. Bars show mean *±* s.d. across mice; mouse labels indicate individual values. Green, blue and purple denote the first, second and last lick-ratio terciles, respectively.Brackets show pairwise mean differences evaluated under exact regrouping of the nine mice into ordered groups of three (1,680 assignments; two-sided tests): first versus second tercile, *p*_regroup_ = 0.1036; first versus last tercile, *p*_regroup_ = 0.9690; and second versus last tercile, *p*_regroup_ = 0.0881. **M:** Relationship between anticipatory licking (*x* axis) and the percentage of cells classified as neither position-stable nor reward-aligned across the reward-location switch (“other” cells, *y* axis). Points represent individual mice, are labelled by mouse identity and are coloured by lick-ratio tercile; horizontal bars show tercile means. The dashed line shows the weighted linear-regression fit (see Methods). There was no evidence that the fitted slope differed from zero (*β* = *−*0.37, 95% bootstrap CI [*−*2.21, 0.96]; two-sided *p*_boot_ = 0.379 from 3,000 weighted mouse-level bootstrap resamples; exact mouse-label permutation test: two-sided *p*_perm_ = 0.8403; *n* = 9 mice; 362,880 permutations). The ordered-tercile statistic was *T* = 0.67 (exact ordered-tercile permutation test: two-sided *p*_tercile_ = 0.825, one-sided *p*_tercile_ = 0.413 for *T >* 0; 1,680 ordered 3/3/3 assignments). **N**: Relationship between anticipatory licking (*x* axis) and the density of selected, previously reward-aligned cells whose post-switch reward-aligned peak fell within the current reward vicinity (*y* axis). Points represent individual mice, are labelled by mouse identity and are coloured by lick-ratio tercile; horizontal bars show tercile means. The dashed line shows the weighted linear-regression fit. The fitted slope was positive (*β* = 0.24, 95% bootstrap CI [0.11, 0.34]; one-sided *p*_boot_ = 0.005 for *β >* 0 from 3,000 weighted mouse-level bootstrap resamples; exact mouse-label permutation test: two-sided *p*_perm_ = 0.0672, one-sided *p*_perm_ = 0.0436 for *β >* 0; *n* = 9 mice; 362,880 permutations). **O**: Relationship between anticipatory licking (*x* axis) and the density of selected, previously reward-aligned cells whose post-switch reward-aligned peak remained within the previous reward zone (*y* axis). Plotting conventions and statistical procedures are as in panel N. The fitted slope was negative (*β* = *−*0.12, 95% bootstrap CI [*−*0.15*, −*0.06]; one-sided *p*_boot_ = 0.007 for *β <* 0 from 3,000 weighted mouse-level bootstrap resamples; exact mouse-label permutation test: two-sided *p*_perm_ = 0.0505, one-sided *p*_perm_ = 0.0240 for *β <* 0; *n* = 9 mice; 362,880 permutations).

**Fig. S7.**
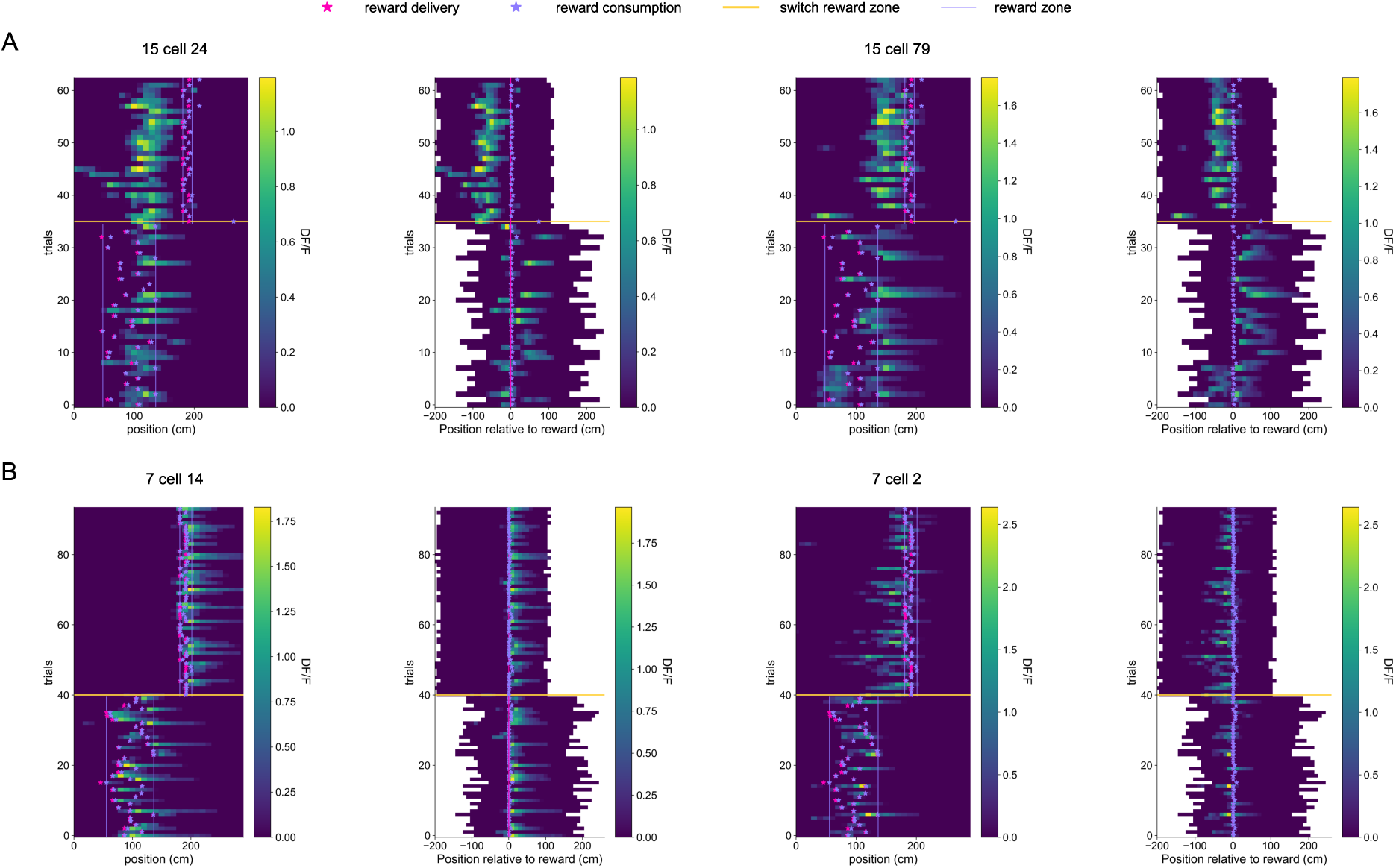
Examples of distinct reward-related remapping patterns across the reward-location switch. **A:** Two example place cells (cells 24 and 79) from reward-responsive mouse J15 that were reward-aligned before the switch and remained active near the previous reward location after the switch. For each cell, trial-by-trial activity is shown in the position reference frame (left) and in reward-consumption aligned coordinates (right). Heat-map colour indicates Δ*F/F*, and trials are ordered chronologically from bottom to top. The horizontal yellow line marks the switch between the pre- and post-switch reward locations. In the position-reference-frame plots, magenta stars indicate reward delivery, lavender stars indicate reward consumption, and paired lavender vertical lines delimit the range of reward-consumption locations within each switch phase. In the reward-consumption-aligned plots, reward consumption defines 0 cm; magenta stars indicate reward-delivery positions relative to reward consumption. **B:** Two example place cells (cells 14 and 2) from reward-anticipatory mouse J7 that were reward-aligned before the switch and shifted their activity with the reward after the switch. Cells and plotting conventions are as in **A**.

## Supplementary Methods

### Suite2p processing parameters

Two-photon imaging time-series movies were imported into Suite2p for rigid and non-rigid motion correction. Putative cell regions of interest were extracted from motion-corrected movies using the parameters below.

### Peak-location analysis

The position of maximum activity was defined as the 10 cm bin in which average activity was greatest. For proximal–distal comparisons, average activity was computed separately for proximal and distal reward trials (Fig. 2 and Supplementary Figs. S3 and S4). For switch analyses, average activity was computed separately before and after the reward-location switch (Fig. 3 and Supplementary Figs. S5 and S6). For proximal–distal pooled heatmaps, cells were included when they met the place-cell criterion before the switch; for across-switch pooled heatmaps, cells were included when they met the place-cell criterion both before and after the switch. Activity maps were normalized by each cell’s maximum activity. Cells were ordered by the peak location of the even-trial map from the ordering condition; displayed activity was computed from the odd-trial map for independently ordered maps and from the corresponding comparison trials when distal- or after-switch activity was displayed in the proximal- or before-switch order (Figs. 2 and 3; Supplementary Figs. S3 and S5).

**Table S1.**
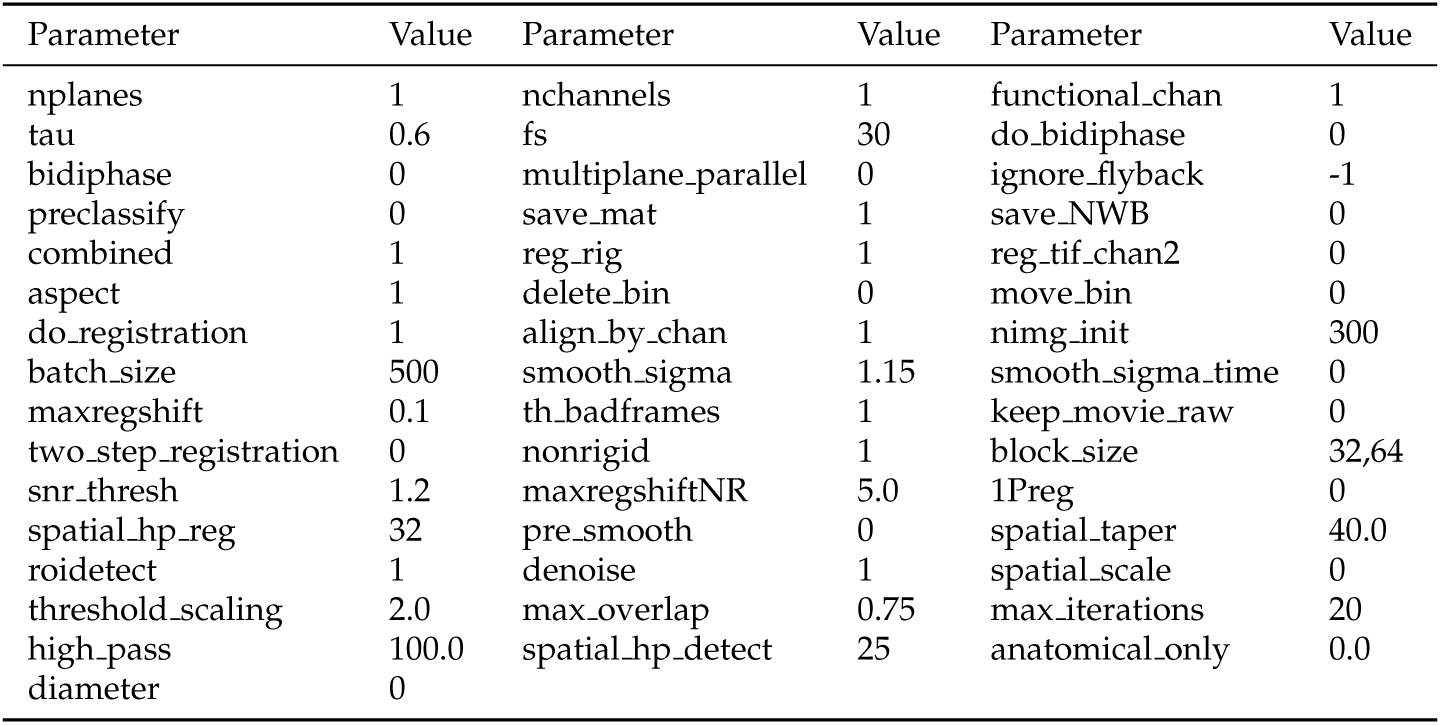
Suite2p parameters.

For two-dimensional scatter peak-location visualizations, the *x−* (showing the argmaximum before the switch or on proximal trials) and *y−* (showing the argmaximum after or on distal trials) coordinates, were fit with the `gaussian kde` function from `scipy.stats`, which estimates the probability density function of a random variable non-parametrically. Heatmaps show these Gaussian kernel density estimates. Dots were slightly jittered to visualise density at equal *x* and *y* coordinates (Fig. 2, Fig. 3, Supplementary Fig. S3 and Supplementary Fig. S5).

### Cell percentages and percentages per centimeter

Cell percentages were computed as the fraction of cells satisfying the relevant criterion relative to the analyzed place-cell population for that comparison. Percentages per centimeter were computed by dividing the percentage of cells in a spatial region by the length of that region in centimeters, allowing comparisons between regions of different widths.

### Mouse-level and shuffle analyses

Mouse-level regressions relating anticipatory licking to neural measures used one point per mouse (Fig. 2D, Fig. 3D and Supplementary Figs. S4D and S6D,M–O). The behavioural value for each mouse was the median adjusted pre/(pre+post) lick ratio across included trials in the analysed phase. For normalized lick-ratio analyses, this value was centred by the across-mouse median and divided by the median absolute deviation.

Weighted least-squares regressions were fit as *y* = *β*_0_ + *β*_1_*x*, where *x* was the mouse-level median adjusted pre to post lick ratio value and *y* was the mouse-level neural measure. For cell-class percentage analyses, *y* was the percentage of analysed cells in the relevant class. The default mouse weight was

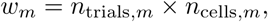

where *n*_trials*,m*_ was the number of included pre-switch behavioural trials and *n*_cells*,m*_ was the number of analysed cells for mouse *m*. Cell-count-only and equal-weight regressions were generated as sensitivity analyses where indicated. Weighted slopes were obtained by solving the normal equations

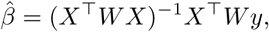

with *X* containing an intercept and the behavioural predictor. The reported slope *β*_1_ therefore represents the change in neural percentage per unit change in the plotted behavioural variable. For normalized behavioural predictors, slopes are expressed per median-absolute-deviation unit of the lick-ratio measure.

Bootstrap confidence intervals for slopes were computed from 3,000 bootstrap resamples of mice for the mouse-level regression analyses (Fig. 2D, Fig. 3D,H and Supplementary Figs. S4D and S6D,H,M–O). For weighted regressions, mice were sampled with replacement with probability proportional to their regression weights, and the weighted slope was recomputed for each bootstrap sample. The reported 95% confidence interval was the 2.5^th^ to 97.5^th^ percentile of the bootstrap slope distribution. Directional bootstrap *p* values were computed as the fraction of bootstrap slopes crossing zero in the direction opposite to the prespecified alternative, with a one-count correction. Prespecified alternatives were positive slopes for reward-referenced cell measures, negative slopes for position-stable cell measures and positive slopes for before-versus-after stability measures. Bias measures used two-sided tests.

Mouse-label permutation *p* values for regression slopes were computed by holding the behavioural values and weights fixed and permuting the neural values across mice. For nine-mouse analyses in Fig. 2D, Fig. 3D,H and Supplementary Figs. S4D and S6D,H,M–O, exact tests enumerated all 9! assignments. Two-sided *p* values were the fraction of permuted slopes with absolute value at least as large as the observed absolute slope. One-sided *p* values were the fraction of permuted slopes at least as large as the observed slope for positive alternatives, or at most as small as the observed slope for negative alternatives. For analyses reported with Monte Carlo rather than exhaustive permutation, 10,000 random permutations were used and *p* values were computed as (*k* + 1)*/*(*N* + 1), where *k* was the number of permuted slopes at least as extreme as the observed slope and *N* was the number of valid permutations. This Monte Carlo procedure was used only for regression or regrouping controls where exhaustive enumeration was not used; the main nine-mouse weighted slope tests and tercile tests used exact enumeration.

For tercile analyses, mice were ordered by median adjusted pre/(pre+post) lick ratio and assigned to low, middle and high groups of three mice each (Fig. 2D, Fig. 3D,I,J and Supplementary Figs. S4D and S6D,L–O). The ordered-tercile statistic was the ordinary least-squares slope of the mouse-level neural value against tercile codes 0, 1 and 2. This tercile statistic was unweighted because the inference unit was the mouse and each tercile contained the same number of mice. Exact *p* values were computed over all ordered assignments of nine mice into three groups of three: three low-tercile mice, three middle-tercile mice from the remaining six and the remaining three high-tercile mice, giving 1,680 assignments. Two-sided *p* values used the absolute ordered-tercile statistic; one-sided *p* values used the observed direction of the statistic. When weighted continuous regression lines were shown together with tercile summaries, the continuous regression used the weighted least-squares procedure described above, whereas tercile group means and ordered-tercile tests remained mouse-level and unweighted.

For analyses of post-switch peak locations of previously reward-aligned cells, cells were selected solely from their pre-switch activity by requiring a reward-consumption-aligned peak within the reward window (*−*20 to +30 cm; Fig. 3I,J and Supplementary Fig. S6I–O). Post-switch peaks were then classified as falling in the current reward zone, the previous reward zone or the remaining track. Region occupancy was summarized as percentage of selected cells per centimetre, computed as 100*×* the number of cells in a region divided by the product of the number of selected cells and the width of that region in centimetres. Within each lick-ratio tercile, paired comparisons between current, previous and remaining-track densities used exact sign-flip tests across mice. Across-tercile comparisons and pairwise contrasts between tercile means were evaluated under all 1,680 exact assignments of the nine mice to three ordered groups of three. For each pairwise contrast, the test statistic was the difference between the corresponding two group means within each 3/3/3 assignment. Pooled zone-versus-remaining-track comparisons, where reported, used one-sided proportion z-tests on region counts normalized by spatial width.

For shuffle-enrichment analyses, post-switch peak locations were permuted within mouse while pre-switch peak locations and mouse identity were held fixed (Fig. 3 and Supplementary Fig. S3). For each mouse and cell-class criterion, 10,000 shuffles were used to estimate the null fraction of cells satisfying that criterion. Enrichment was defined as the observed fraction minus the mean shuffled fraction. Mouse-level shuffle *p* values were computed as (*k* + 1)*/*(*N* + 1), where *k* was the number of shuffled fractions greater than or equal to the observed fraction and *N* = 10,000. Regressions of shuffle enrichment against behavioural measures used mouse-level regression and permutation procedures as described above.

## References

[1] Radvansky, B. A., Oh, J. Y., Climer, J. R. & Dombeck, D. A. Behavior determines the hippocampal spatial mapping of a multisensory environment. Cell reports 36 (2021).

[2] Sanders, H., Wilson, M. A. & Gershman, S. J. Hippocampal remapping as hidden state inference. Elife 9, e51140 (2020).

[3] O’Keefe, J. & Dostrovsky, J. The hippocampus as a spatial map: preliminary evidence from unit activity in the freely-moving rat. Brain research (1971).

[4] Leutgeb, J. K. et al. Progressive transformation of hippocampal neuronal representations in “morphed” environments. Neuron 48, 345–358 (2005).

[5] Muller, R. U. & Kubie, J. L. The effects of changes in the environment on the spatial firing of hippocampal complex-spike cells. Journal of Neuroscience 7, 1951–1968 (1987).

[6] Gauthier, J. L. & Tank, D. W. A dedicated population for reward coding in the hippocampus. Neuron 99, 179–193 (2018).

[7] Qian, F. K., Li, Y. & Magee, J. C. Mechanisms of experience-dependent place-cell referencing in hippocampal area ca1. Nature neuroscience 28, 1486–1496 (2025).

[8] McKenzie, S. et al. Hippocampal representation of related and opposing memories develop within distinct, hierarchically organized neural schemas. Neuron 83, 202–215 (2014).

[9] Sosa, M., Plitt, M. H. & Giocomo, L. M. A flexible hippocampal population code for experience relative to reward. Nature Neuroscience 28, 1497–1509 (2025).

[10] Tessereau, C., Xuan, F., Jack, R. M., Dayan, P. & Dombeck, D. Navigating uncertainty: reward location variability induces reorganization of hippocampal spatial representations. bioRxiv (2025).

[11] Pettit, N. L., Yuan, X. C. & Harvey, C. D. Hippocampal place codes are gated by behavioral engagement. Nature neuroscience 25, 561–566 (2022).

[12] Krishnan, S., Heer, C., Cherian, C. & Sheffield, M. E. Reward expectation extinction restructures and degrades ca1 spatial maps through loss of a dopaminergic reward proximity signal. Nature communications 13, 6662 (2022).

[13] Barnstedt, O., Mocellin, P. & Remy, S. A hippocampus-accumbens code guides goal-directed appetitive behavior. Nature Communications 15, 3196 (2024).

[14] Kentros, C. G., Agnihotri, N. T., Streater, S., Hawkins, R. D. & Kandel, E. R. Increased attention to spatial context increases both place field stability and spatial memory. Neuron 42, 283–295 (2004).

[15] Packard, M. G. & McGaugh, J. L. Inactivation of hippocampus or caudate nucleus with lidocaine differentially affects expression of place and response learning. Neurobiology of learning and memory 65, 65–72 (1996).

[16] Chersi, F. & Burgess, N. The cognitive architecture of spatial navigation: hippocampal and striatal contributions. Neuron 88, 64–77 (2015).

[17] Goodroe, S. C., Starnes, J. & Brown, T. I. The complex nature of hippocampal-striatal interactions in spatial navigation. Frontiers in human neuroscience 12, 250 (2018).

[18] Zhang, Y. et al. Fast and sensitive gcamp calcium indicators for imaging neural populations. Nature 615, 884–891 (2023).

[19] Dombeck, D. A., Harvey, C. D., Tian, L., Looger, L. L. & Tank, D. W. Functional imaging of hippocampal place cells at cellular resolution during virtual navigation. Nature neuroscience 13, 1433–1440 (2010).

[20] Sheffield, M. E., Adoff, M. D. & Dombeck, D. A. Increased prevalence of calcium transients across the dendritic arbor during place field formation. Neuron 96, 490–504 (2017).

[21] Pachitariu, M., et al. Suite2p: beyond 10,000 neurons with standard two-photon microscopy. bioRxiv (2017).

[22] Climer, J. R. & Dombeck, D. A. Information Theoretic Approaches to Deciphering the Neural Code with Functional Fluorescence Imaging. eNeuro 8 (2021).

